# Stereo-specific Lasofoxifene Derivatives Reveal the Interplay between Estrogen Receptor Alpha Stability and Antagonistic Activity in ESR1 Mutant Breast Cancer Cells

**DOI:** 10.1101/2021.08.27.457901

**Authors:** David J. Hosfield, Sandra Weber, Nan-Sheng Li, Madline Sauvage, Emily Sullivan, Estelle Nduwke, Ross Han, Sydney Cush, Muriel Lainé, Sylvie Mader, Geoffrey L. Greene, Sean W. Fanning

## Abstract

Chemical manipulation of estrogen receptor alpha ligand binding domain structural mobility tunes receptor lifetime and influences breast cancer therapeutic activities. Selective estrogen receptor modulators (SERMs) extend ERα cellular lifetime, accumulation, and are antagonists in the breast and agonists in the uterine epithelium and/or in bone. Selective estrogen receptor degraders (SERDs) reduce ERα cellular lifetime/accumulation and are pure antagonists. Activating somatic *ESR1* mutations Y537S and D538G enable resistance to first-line endocrine therapies. SERDs have shown significant activities in *ESR1* mutant setting while few SERMs have been studied. To understand whether chemical manipulation of ERα cellular lifetime and accumulation influences antagonistic activity, we synthesized a series of methylpyrollidine lasofoxifene derivatives that maintained the drug’s antagonistic activities while uniquely tuning ERα cellular accumulation. These molecules were examined alongside a panel of antiestrogens in live cell assays of ERα cellular accumulation, lifetime, SUMOylation, and transcriptional antagonism. High-resolution x-ray crystal structures of WT and Y537S ERα ligand binding domain in complex with the methylated lasofoxifene derivatives, SERMs, and SERDs show that molecules that favor a highly buried helix 12 conformation achieve the greatest transcriptional suppression activities. Together these results show that chemical reduction of ERα cellular lifetime does not necessarily correlate with transcriptional antagonism in *ESR1* mutated breast cancer cells. Importantly, our approach shows how minor chemical additions modulate receptor cellular lifetime while maintaining other activities to achieve desired SERM or SERD profiles.

**SIGNIFICANCE:** This study shows that antiestrogens that enforce a wild-type-like antagonist conformation demonstrate improved therapeutic activities in hormone-resistant breast cancer cells harboring Y537S and D538G *ESR1*.

## INTRODUCTION

The pro-oncogenic cellular activities of estrogen receptor alpha (ERα) drive breast cancer pathogenesis. ERα is overexpressed in approximately 70% of breast cancers and targeted endocrine therapies are given to prevent primary disease metastasis. Post-menopausal patients primarily receive aromatase inhibitors (AI), which indirectly inhibits ERα by ablating endogenous estrogens, and may be given in combination with a CDK 4/6 inhibitor^1^. Pre-menopausal patients receive tamoxifen, a selective estrogen receptor modulator (SERM), which directly binds to the receptor and reprograms transcription to induce cellular quiescence^2^. Together, these therapeutic paradigms significantly reduce the 5-year risk of recurrence and continually show improved disease outcomes^3, 4^.

Acquired resistance to endocrine therapies remains a major source of breast cancer mortality. Approximately 50% of patients present acquired or *de novo* resistance to endocrine therapies after an average of 5-years^5^. Deep genomic sequencing of endocrine-resistant, ER+, and metastatic breast cancers revealed the presence of *ESR1* ligand binding domain mutations (*ESR1*muts) at a rate of approximately 25%^5–7^. Y537S (14%) and D538G (36%) are the two most prevalent mutations and account for nearly 50% of those identified. Both mutations, but especially Y537S, enable hormone-free transcriptional activity, resistance to inhibition by 4-hydroxytamoxifen (4OHT, the active metabolite of tamoxifen), and are associated with a more aggressive disease phenotype^8^. *ESR1*muts have also been identified in cultured breast cancer cells and become enriched as they become resistant to endocrine therapies, suggesting a latent and selectable mechanism of acquired drug resistance^9^. In addition to mimicking the genomic actions of estradiol (E2) stimulation, *ESR1*muts bind to unique cistromes and promote allele-specific transcriptional programs compared to E2-stimulated WT receptor^10^. Mutation-induced molecular alterations enable resistance to clinically approved antiestrogens and a more aggressive metastatic progressive disease.

Biochemical studies and x-ray crystal structures have shown that both mutations favor an E2-like agonist conformation in the absence of hormone^11, 12^. This favored agonist conformation reduces the binding affinity of ERα ligands and alters the therapeutically important antagonist receptor conformation of the selective estrogen receptor modulator (SERM) 4OHT^11^. Competitive antiestrogens with selective estrogen receptor degrader/downregulator (SERD) activities with improved potencies show superior therapeutic antagonistic activities compared to 4OHT in breast cancer cells harboring Y537S or D538G *ESR1*^13^. The SERD Fulvestrant, but not 4OHT, completely ablated Y537S ERα transcriptional activity in breast cancer cells^5^. Fulvestrant’s activity stems from its ability to antagonize ERα transcription and reduce its cellular lifetime (degrade/downregulate receptor) by inducing post-translational modifications including ubiquitination, SUMOylation, and proteasomal degradation^14–16^. Tamoxifen and other selective estrogen receptor modulators (SERMs) also antagonize transcription but induce a stable antagonist conformation that enhances ERα nuclear lifetime^13^. SERMs show partial agonist activities in the uterine endometrium and/or in bone while SERDs are pure antagonists ^17^. Because fulvestrant possesses unfavorable pharmacological properties, especially poor solubility and lack of oral bioavailability, large-scale efforts brought about the development and clinical evaluation of orally available SERDs^18–26^. These molecules possess functional groups including carboxylic acids, azeditines, and pyrrolidines that favor the H12 antagonist conformation and induce proteasomal degradation by increasing its conformational mobility, although their impact on receptor post-translational modifications remains uncharacterized^17, 18, 26, 27^.

Recent studies suggest that ERα degrading activities may not be required for therapeutic efficacy in breast cancer cells with recurring hotspot *ESR1* mutations. Next generation antiestrogens with mixed SERM/SERD activities including bazedoxifene and lasofoxifene, which possess weaker ERα degrading/downregulating activities, show potent anti-cancer activities in breast cancers harboring *ESR1*muts ^13, 28–30^. In addition, mutations abolishing induction of receptor SUMOylation reduced but did not abolish transcriptional suppression by fulvestrant in breast cancer cells^16^. Further, in the WT *ESR1* setting, Wardell et al. have shown that antagonism drives fulvestrant’s anti-cancer activities even when degradation is abolished or incomplete^31, 32^. Ectopic overexpression of ERα superseded the proteasome’s ability to degrade receptor but fulvestrant retained transcriptional inhibition activity^32^. Further, a recent re-evaluation of fulvestrant’s pharmacologic and pharmacokinetic properties suggests that its efficacy of transcriptional antagonism, rather than ERα degrading activities, primarily drives anti-tumoral response in different models of ER+ breast cancer^31^. Therefore, therapeutic efficacy may depend on other aspects of ERα antagonism besides degrading/downregulating activities. In light of this recent work, the role of ERα modification by ubiquitin and SUMO, cellular lifetime, and accumulation in anti-cancer therapeutic activities, especially with the *ESR1_muts_*, remains unclear.

In this study, we synthesized and characterized a series of methyl derivatives of the SERM/SERD lasofoxifene that uniquely enhanced or diminished WT and mutant ERα cellular accumulation without sacrificing potency to study the role of receptor stability on transcriptional antagonistic and antiproliferative activities. We evaluated these molecules against a panel of SERMs and SERDs (**Figure 1**) that represent a comprehensive spectrum of clinical and preclinical antiestrogenic small molecules. We developed a novel high-throughput live-cell approach to quantify and observe the kinetics of ERα cellular accumulation within living breast cancer cells upon head-to-head treatment with SERM or SERD. Because fulvestrant-induced SUMOylation of ERα correlates with shortened kinetics of interaction with DNA, lowered accessibility of estrogen response elements, and more efficient repression of ERα target genes in the presence of fulvestrant^15, 16^, bioluminescence resonance energy transfer (BRET) was used to examine differences between SERM and SERD-induced SUMOylation for WT, Y537S, and D538G ERα. Reporter gene assays quantifying the impact of SERM and SERD treatment on WT and heterozygous Y537S/WT and D538G/WT transcriptional antagonism in MCF7 breast cancer cells revealed that antiestrogen efficacy at suppressing transcription did not correlate with reduced receptor accumulation.

**Figure 1:**
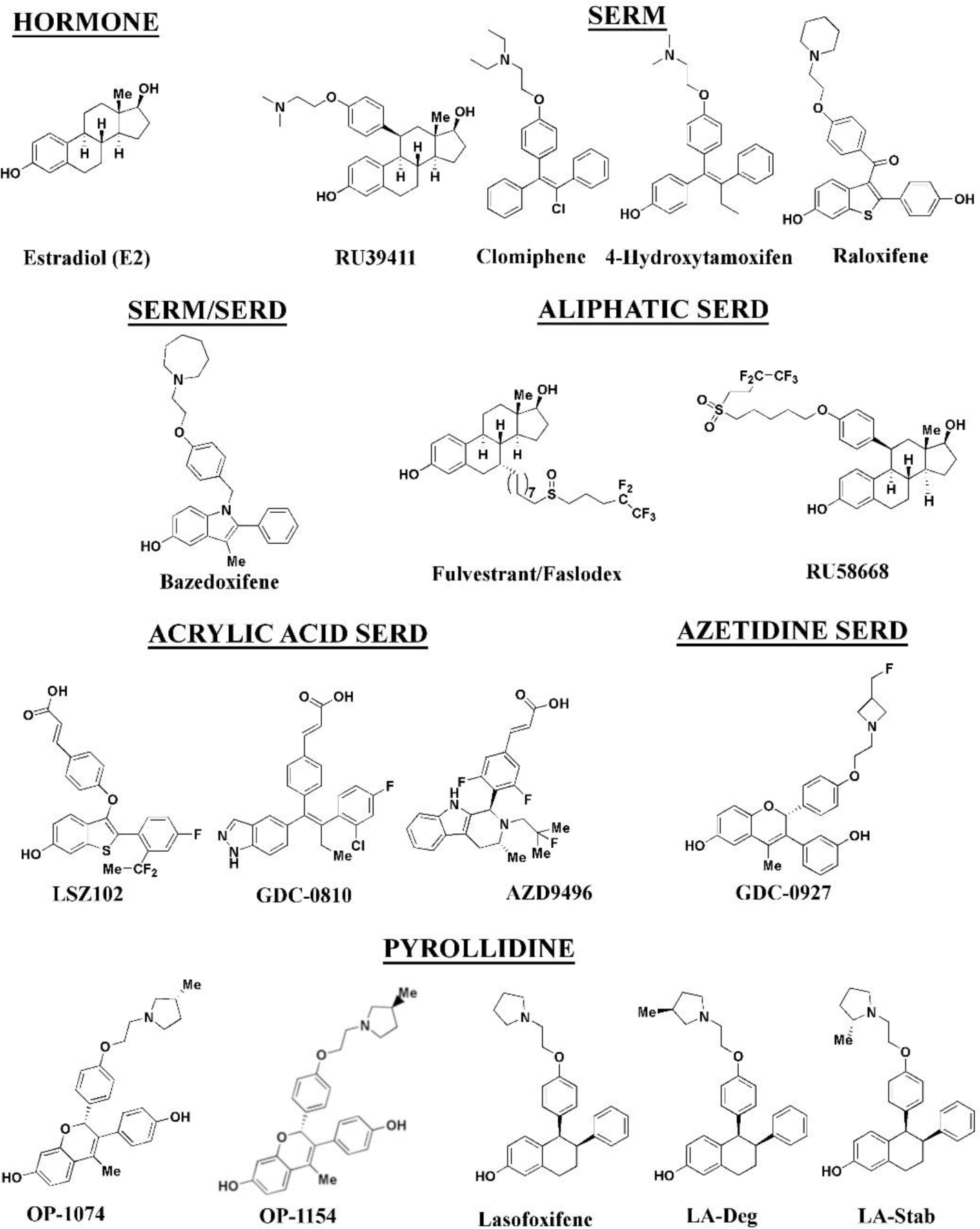
Estrogen receptor alpha ligands evaluated in this study including estradiol, selective estrogen receptor modulators (SERMs), and selective estrogen receptor degraders/downregulators (SERDs).

Cellular proliferation studies in MCF7 with heterozygous WT/Y537S *ESR1*, which possesses the greatest degree of antiestrogen-resistance, show that manipulating ERα stability did not enhance anti-proliferative activities in cultured cells. X-ray crystal structures of Y537S ERα LBD in complex with SERMs and SERDs reveal a conserved ligand-dependent conformational pose that enables marked transcriptional antagonistic activities for *ESR1* mutant.

## RESULTS

### Ligand Influence on ERα Expression in Live Breast Cancer Cells

We developed a high-throughput assay to quantify differences in ERα expression and cellular lifetime based on ligand or mutation within live T47D breast cancer cells engineered to express a doxycycline (dox) inducible halo-tagged ERα, Populations that stably incorporated the receptor were enriched by flow-sorting. The cell permeable halo-618 fluorophore was used to follow ERα expression and cellular lifetime. To determine ERα accumulation, cells were distributed into 96-well plates in estrogen depleted medium followed by simultaneous addition of Dox, halo-618, and hormone or antiestrogen for 24-hours. Using a range of dox concentrations, we found that 1 μg/mL was the lowest dose needed to achieve maximal signal-to-noise ratio (**Figure 2-figure supplement 1**). To measure IC_50_ of ERα stabilization or degradation/downregulation, cells were simultaneously treated dox, halo-618 and hormone (estradiol [E2]), SERMs, SERM/SERDs, or SERDs at concentrations varying between 0.1 nM and 5 μM for 24-hours. **Figure 2** shows representative curves experiments shown as the mean of two biological replicates ± standard deviation after normalization to cell count by phase contrast. **Figure 2-supplemental file 2** shows IC_50_, R^2^ of fit, and fluorescence normalized to cell count at maximum dose (5 μM) for each ligand and ERα mutant. SERMs increase ERα levels in the nucleus, while SERDs decrease ERα by Western-blot^13^. In our assay, SERMs, such as 4-hydroxytamoxifen (4OHT), show a normalized fluorescence greater than 1 at 5 μM, lasofoxifene was neutral between 0.8 and 1, and SERDs were less than 0.8. As expected, the hormone estradiol (E2) induced ERα degradation, but at 0.68 ± 0.05 was approximately 2-3-fold less than the SERDs with the greatest reduction at 0.2 ± 0.03, 0.29 ± 0.04, and 0.33 ± 0.04 for RU58668, fulvestrant (ICI182,870]), and GDC-0927 respectively. The SERMs 4OHT and RU39411 significantly increased ERα signal with a normalized fluorescence of 2.23 + 0.24 and 2.11 + 0.36 at 5 μM treatment respectively. Overall, GDC-0927, fulvestrant, and pipendoxifene (PIP) showed the most potent IC_50_ values at 0.13 ± 0.03, 0.4 ± 0.02, and 0.77 ± 0.03 nM. Notably, many oral SERDs were not significantly different from PIP. To validate our data, we treated normal T47D cells with increasing doses of fulvestrant then used a capillary-based western blotting technique to observe differences in ERα levels after 24-hours normalized to β-actin. We derived an IC_50_ value at 0.34 ± 0.10 with an R^2^ of 0.94 using this method, which is in close agreement with the results from the engineered HT-ERα expressing cells l (**Figure 2-figure supplement 3**).

**Figure 2:**
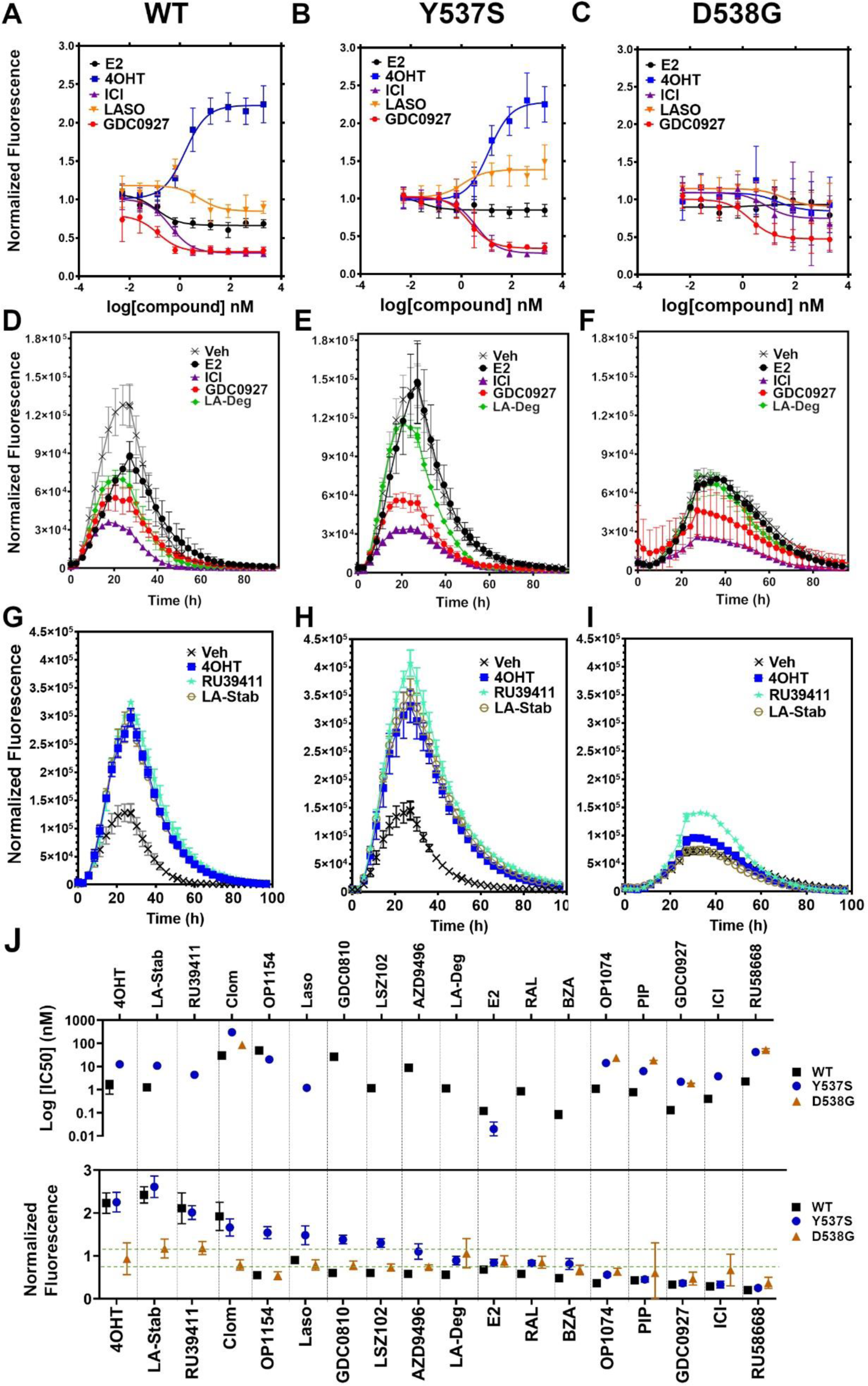
Impact of Ligand and Mutation on Estrogen Receptor Alpha Lifetime in T47D breast Cancer Cells. A-C) Dose-response curves of select ligands after 24-hours for WT (A), Y537S (B), and D538G (C). D-F) Kinetics of ERα cellular lifetime with representative SERDs for WT (D), Y537S (E), and 538G (F). G-I) Kinetics of ERα cellular lifetime with representative SERMs for WT (G), Y537S (H), and 538G (I). J) Summary of IC_50_ and normalized fluorescence at 5 μM compound after 24 hours treatment. Compounds are ordered from left to right based on their maximum normalized fluorescence at the highest dose after 24 hours. Poor IC_50_ fits were omitted. Data above the dashed green lines are classified as SERMs and below are SERDs. Data are shown as the mean of two biological replicates ± standard deviation. All data are normalized to cell count in their respective wells.

### Impact of Y537S and D538G Mutations on ERα Stabilization and Degradation

To understand how Y537S and D538G ERα mutations impact the stabilizing or degrading activities of antiestrogens, we generated stable T47D breast cancer cells that possess dox-inducible halo-tagged Y537S and D538G ERα. SERM-like molecules appeared to retain their abilities to increase receptor nuclear lifetime in the Y537S, but did so with approximately 10-fold reduced IC_50_ values (**Figure 2J** and **Figure 2-supplemental file 2**). Lasofoxifene, which was essentially neutral to slightly SERD-like in the WT cells, showed a SERM-like profile at 1.48 ± 0.22 at 5 μM with Y537S. Interestingly, SERD molecules that possessed carboxylic acids (AZD9496, LSZ102, and GDC0801) all showed more SERM-like profiles with their normalized fluorescence above 1 at 5 μM. Fulvestrant, RU58668, GDC-0927, and PIP were the only molecules that did not show statistically significant differences to their maximal normalized fluorescence with Y537S. However, they did show about a 10-fold decreased IC_50_ compared to WT. Surprisingly, introduction of the D538G mutation showed substantial differences. Notably, impacts of E2 and 4OHT on receptor levels were abolished. Nearly every other SERM and SERD had decreased IC_50_ values and required maximum dosages to achieve an effect on ERα levels. GDC-0927 was the only SERD that attained maximal effect on normalized fluorescence with an IC_50_ < 10 nM at 1.88 ± 0.04 nM.

### Impact of Ligands and Mutations on ERα Cellular Lifetime

One advantage of the halo-ERα live-cell system is that protein lifetime can easily and rapidly be followed in living cells. To uncover whether ligands and activating somatic mutations influenced the cellular lifetime of ERα, we simultaneously induced halo-ERα expression, treated with 1 μM ligand, and measured fluorescence normalized to cell count every four hours for 100 hours. For WT cells, ERα reached a maximum at approximately 24-hours then fell to baseline by 80 hours (**Figure 2D**). Treatment with E2 leads to a reduced expression but the signal converged with vehicle by around 30 hours and had fully returned to baseline around 80 hours. Treatment with SERDs further reduced the signal and led to a rapid return to baseline, at around 45 hours for fulvestrant. Treatment with SERMs greatly enhanced ERα levels over vehicle and extended the time to baseline to past 80 hours. Interestingly, veh-treated WT and Y537S cells showed similar profiles, but the Y537S took slightly longer to reach baseline. Unlike the E2-treated WT cells, E2-treated Y537S cells showed an identical pattern to veh-treated Y537S cells (**Figure 2E**). Surprisingly, Y537S mutation attenuated the rapid degradation of fulvestrant, which extended past 60 hours compared to 40 hours for WT (**Figure 2H**). Unexpectedly, the D538G mutant not only reduced the magnitude of ERα expression, but also the kinetics of its cellular turnover with a greater time to baseline for each SERD (**Figure 2F**). For SERMs, each molecule showed similar profiles of enhancing ERα lifetime in the WT T47D cells (**Figure 2G**). In the 537S cells, they further enhanced stability compared to WT (**Figure 2H**). Surprisingly, RU39411 was the only SERM to elicit any kind of difference on D538G by showing a slight increase in stability, however it was still approximately 2.5-fold less than in the Y537S (**Figure 2I**). These data suggest that, in addition to hormone independent transcriptional activities, both Y537S and D538G mutations uniquely affect ERα cellular lifetime within the T47D breast cancer cell.

### Stereo-Specific Addition of Methyl Groups to Lasofoxifene’s Pyrrolidine Tunes ERα Cellular Lifetime

A major goal of this study was to understand whether chemical manipulation of WT and mutant ERα cellular lifetime enhanced transcriptional antagonistic potencies. Lasofoxifene did not significantly affect ERα signal compared to vehicle across all concentrations. Notably, lasofoxifene possesses a pyrrolidine moiety, similar to OP-1074, which induces ERα degradation through a 3R-methylpyrrolidine^26^. Hence, we synthesized a series of lasofoxifene derivatives with stereo-specific methyl groups at the 2 and 3 positions on the pyrrolidine (**Figure 3A**) We first examined the abilities of the molecules to influence WT halo-ERα lifetime in our engineered T47D cells (**Figure 3B**). LA-3 possesses a 3R methylpyrrolidine (similar to OP1074) and showed a SERD-like profile with an IC_50_ of 26.94 ± 0.4 nM and maximum normalized fluorescence of 0.48 ± 0.07 at 5 μM. LA-5 possesses a 2S methylpyrrolidine and showed a SERM-like profile with an IC_50_ of 15.68 + 0.27 nM and a normalized fluorescence of 1.706 + 0.09 at 5 μM. LA2 and LA4 appeared similar to lasofoxifene in that they did not affect receptor lifetime.

**Figure 3:**
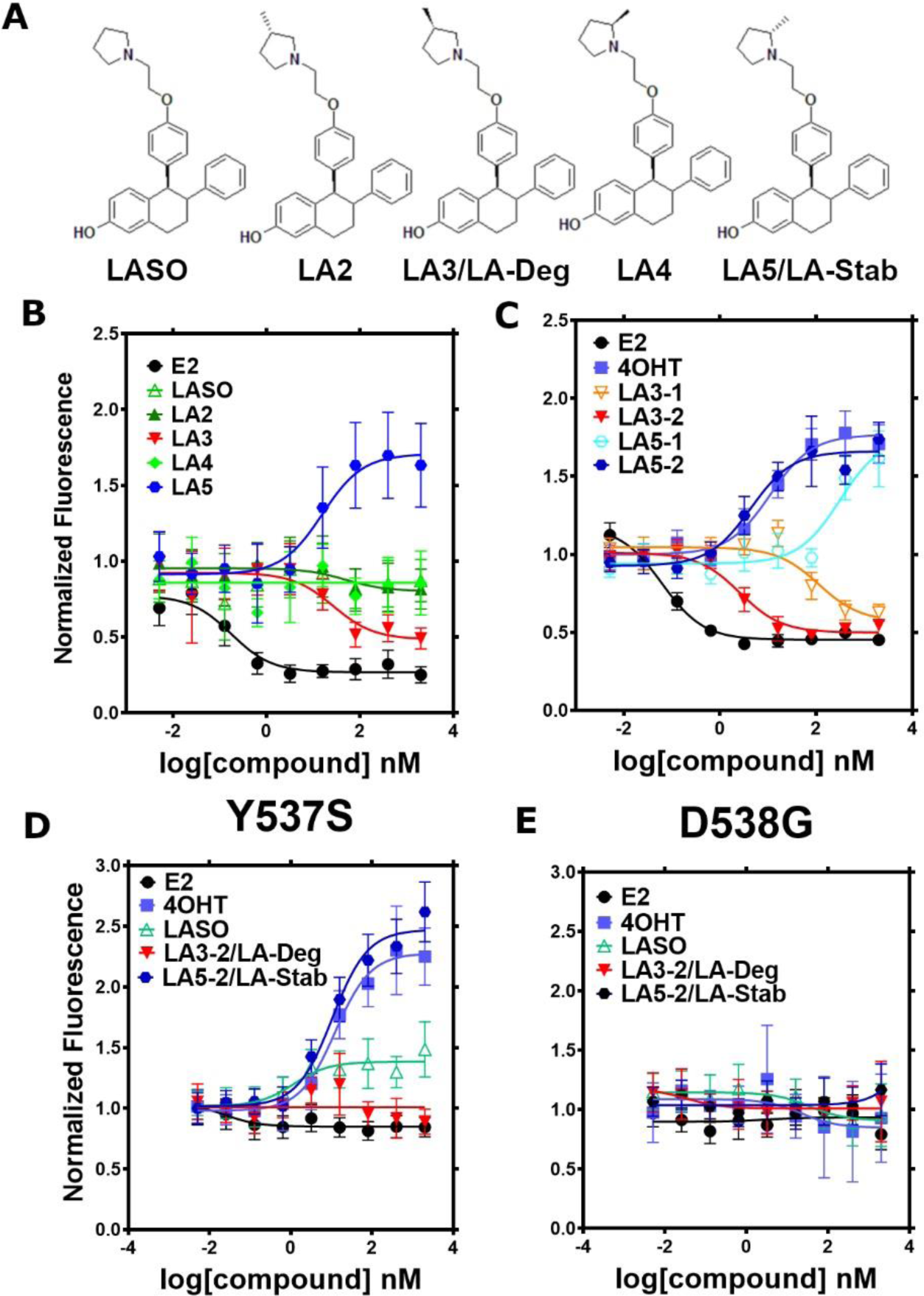
Stereo-specific methyl additions onto the pyrrolidine of lasofoxifene impact ERα levels in T47D breast cancer cells. A) Chemical structures of lasofoxifene and the synthesized stereo-specific methyl derivatives. B) Dose-response curves of hormone (E2) alongside lasofoxifene (LASO) after 24-hour treatment for WT halo-ERα. C) Dose-response curves of chirally purified LA3 and LA5 alongside E2 and 4-hydroxytamoxifen (4OHT) for WT halo-ERα. LA3/5-1 and LA3/5-2 represent the first and second major peaks separated by chiral affinity chromatography respectively. D/E) Dose-response curves of LA-Deg and LA-Stab for Y537S and D537G halo-ERα after 24 hours compared to E2, LASO, and 4-hydroxytamoxifen (4OHT). Data are shown as the mean of two biological replicates ± standard deviation.

LA3 and LA5 were synthesized as racemic mixtures. To understand whether one chiral species accounted for most of the activity, we performed chiral separation and were able to resolve two major peaks (**Figure 3-figure supplement 1**). For both LA3 and LA5, we measured significantly improved IC_50_ values for the second peak versus the first peak (**Figure 3C**). For LA3 peak 1 (LA3-1) and LA3-2 the IC_50_ was 112.9 ± 0.23 nM and 2.58 ± 0.13 nM respectively. The normalized fluorescent signals at 5 μM were somewhat decreased for LA3-1 versus LA3-2 at 0.57 + 0.02 and 0.49 + 0.02 respectively. They were also slightly decreased at 1.75 + 0.02 and 1.66 + 0.04 for LA5-1 and LA5-2 respectively. As such, chiral separation substantially improved the IC_50_ but not the fluorescence at maximum dose. LA3-2 and LA5-2 are subsequently referred to as LA-Deg and LA-Stab for Lasofoxifene Degrader and Stabilizer respectively.

We next examined the abilities of LA-Deg and LA-Stab to affect the cellular lifetime of Y537S and D538G halo-ERα in T47D breast cancer cells. In Y537S cells, LA-Stab demonstrated a slightly improved IC_50_ over 4OHT at 10.62 ± 0.12 and 12.42 ± 0.15 nM respectively. LA-Stab also increased Y537S levels to a slightly greater extent than 4OHT at maximum dose at a signal of 2.47 + 0.06 and 2.27 + 0.07 respectively (**Figure 3D**). Interestingly, the Y537S essentially neutralized the SERD activities of LA-Deg with a normalized fluorescence of 1.01 + 0.04 at 5 μM. Laso showed a more SERM-like profile with a normalized fluorescence of 1.38 + 0.03 at 5 μM. As with other molecules, the D538G mutant completely neutralized the LA-Deg and LA-Stab with normalized fluorescent signals around 1 at 5 μM treatment. In the kinetics/lifetime assays, the methylated Laso derivatives showed similar profiles compared to other antagonists (Figure 2D-I). The values for IC_50_ and normalized fluorescence at maximum dose for LA-Deg and LA-Stab are shown alongside other antagonists in **Figure 2J**. Overall, these data show that the methylated Laso derivatives have different impacts on WT ERα accumulation and lifetime but are affected by Y537S and D538G mutations similar to other antiestrogens with comparable properties.

### Impact of Ligands and Mutations on SUMOylation

The activating somatic mutations Y537S and D538G differentially influenced how ligands affect ERα cellular lifetime. Fulvestrant induces both ubiquitination and SUMOylation of ERα in breast cancer cells, impacting its stability and activity^15, 17^. SUMOylation of ERα can be measured in HEK293T cells using bioluminescence resonance energy transfer (BRET) between Renilla Luciferase (RLucII) tagged ERα and YFP tagged SUMO3^16, 33^. **Figure 4** shows representative SUMO BRET response curves for WT, Y537S, and D538G ERα with a panel of antiestrogens. **Figure 4-supplemental file 1** shows IC_50_ values and maximum BRET ratios for the SUMOylation experiments. In both WT and Y537S cells, fulvestrant induces the greatest degree of SUMO3 recruitment, but the IC_50_ values and maximum BRET ratios are decreased with the Y537S mutant. GDC0927 is the next most efficacious antiestrogen and its potency trends similarly to fulvestrant with the Y537S mutant. LA-Deg induces but to a lesser maximum than fulvestrant and GDC0927 in WT and is further reduced in Y537S, while LA-Stab did not induce SUMOylation with either receptor. Interestingly, RAL, a SERM with weak SERD-like activity with WT but not mutant ERα in our cellular lifetime and accumulation assays, also induced low levels of SUMOylation in the WT but not the Y537S cells. Overall, these assays show a correlation between a molecule’s ERα-degrading activities and induction of SUMOylation. However, the acrylic acid SERD AZD9496 did not induce SUMOylation with any ERα construct.

**Figure 4:**
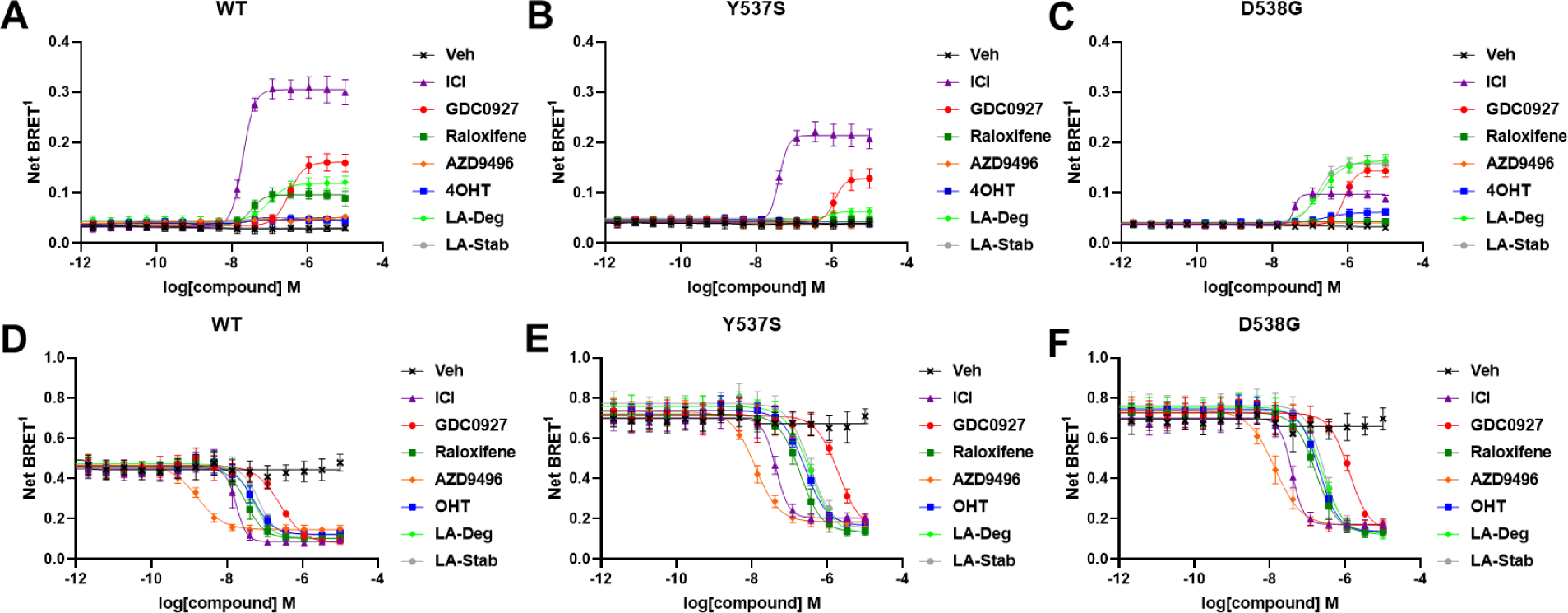
Impact of Y537S and D538G mutation on ligand-induced ERα SUMOylation and SRC1 coactivator binding. SUMOylation of WT (A), Y537S (B), or D538G (C) ERα in the presence of vehicle, fulvestrant, GDC0927, raloxifene, AZD9496, 4-hydroxytamoxifen (4OHT), lasofoxifene-degrader (LA-Deg), or lasofoxifene-stabilizer (LA-Stab). Data are shown as the mean ± s.e.m., *n* = 3 to 5 biological replicates. Association of WT (D), Y537S (E), and D538G (F) ERα and the receptor-interacting domain of SRC1. Data are shown as the mean ± s.e.m., *n* = 3 biologic replicates.

As in our cellular lifetime studies, the D538G mutant showed the greatest alterations in induced SUMOylation compared to WT and Y537S ERα in the BRET assay. Fulvestrant induced about 25% of the SUMOylation for D538G compared to WT ERα. GDC0927 was less affected in its capacity to induce SUMOylation by D538G, albeit its potency was reduced, consistent with our lifetime/accumulation experiments. Surprisingly, LA-Stab and LA-Deg displayed similar activity, contrary to observations with WT and Y537S ERα. Interestingly, 4OHT also induced low levels of SUMOylation in the D538G mutant that were not observed for WT and Y537S. These gains in SUMOylation for LA-Stab and 4OHT correlate with their loss of stabilization activity observed in T47D cells. Because fulvestrant showed significant differences in SUMOylation for the D538G mutant, we examined whether addition of estradiol (E2) further affected its efficacy for induction of this post-translational modification (**Figure 4-figure supplement 2**). Both the efficacy and potency of fulvestrant for SUMOylation were markedly reduced, similar to our previous studies in which they were significantly reduced for the D538G mutant. Together, these studies show that the D538G activating somatic *ESR1* mutant uniquely affects receptor SUMOylation in response to SERDs, in agreement with our measured differences in cellular lifetime.

### Hotspot Mutations Enhance Coactivator Recruitment and Reduce Inhibitory Potencies in Cells

Ligand-dependent transcriptional activities depend on the association of coactivator proteins with ERα to form functional complexes^34^. SERMs and SERDs elicit a structural rearrangement of the ERα LBD to sterically preclude binding of coactivators in the activating function-2 cleft^17^. Mutations affect the ligand-dependence, antagonistic efficacies and potencies of SERMs and SERDs, and specific coactivator proteins that associate with ERα^10, 11, 13^. As such, we used BRET to understand whether SERD molecules showed improved abilities to inhibit the binding of the receptor-interacting domain (RID) of the coactivator SRC1 with WT and mutant ERα in cells. RLucII tagged ERα and YFP tagged SRC1 RID were used to measure differences in association in HEK293T cells. **Figure 4D** shows representative curves for WT and mutant ERα-SRC1 RID binding. **Figure 4-supplemental file 3** shows IC_50_ values and maximum BRET ratios for the coactivator binding experiments. Overall, enhanced basal BRET ratios are observed for the mutants compared to WT ERα. The mutations also decreased the potencies of all SERMs and SERDs to inhibit coactivator binding, consistent with earlier findings for 4OHT and bazedoxifene (BZA) using purified recombinant proteins *in vitro*^13^. Interestingly, all molecules showed similar efficacies in inhibiting this interaction at the greatest concentrations. AZD9496 showed the greatest potency for all ERα species, but is approximately 10-fold reduced in the mutants. Fulvestrant was the next most potent. Raloxifene, 4OHT, LA-Deg, and LA-Stab showed similar potencies in cells with WT and mutant ERα. GDC0927 showed the least potent inhibition of coactivator association in the WT and mutant ERα assays. Because SERDs showed varied potencies in prohibiting coactivator association with the ERα mutants and did not display greater efficacies, we conclude that induction of ERα SUMOylation does not correlate with repression of SRC1 RID recruitment in cells. Importantly, the mutants uniquely affect which coactivator proteins associate with ERα^10^. Therefore, additional studies will be needed to identify preferred antiestrogens to target mutant-specific cofactor interactions.

### SERMs and SERDs Antagonize *ESR1* Mutant Transcriptional Activities

We used reporter gene assays to measure transcriptional antagonistic potencies of antiestrogens in breast cancer cells harboring WT, Y537S, or D538G *ESR1*. A 3x-estrogen response element DNA sequence upstream of a GFP reporter was stably incorporated into MCF7 cells harboring WT, heterozygous WT/Y537S, or heterozygous WT/D538G *ESR1* (donated by Dr. Ben Ho Park). Cells with stable incorporation of the gene cassette were selected for and enriched using flow sorting. Cells were placed in serum-starved media for 48 hours then treated with increasing concentrations of ligand. ERα transcription was determined by quantitating the total integrated green fluorescence of these cells 24 hours later (**Figure 5**, **Figure 5-supplemental file 1**). For WT cells: GDC0927, RAL, and OP1074 showed the most potent transcriptional inhibition at IC_50_ = 0.03 ± 0.08, 0.11 ± 0.08, and 0.14 ± 0.05 nM respectively, compared to 0.35 ± 0.06 for fulvestrant (**Figure 5D and E**). At the 5 μM maximum dose, RU39411, RAL, and BZA showed the greatest reduction in transcription at 0.14 ± 0.02, 0.16 ± 0.02, and 0.17 ± 0.01. In the WT/Y537S heterozygous cells GDC0927, RAL, and lasofoxifene showed the most potent inhibition at IC_50_ = 0.95 ± 0.51, 2.16 ± 0.67, and 2.88 ± 0.34 nM respectively. At the maximum dose: RU39411, BZA, and PIP showed the greatest reduction in transcription at 0.33 ± 0.02, 0.32 ± 0.05, and 0.32 ± 0.04 respectively. As previously reported, no antiestrogen completely reduced transcriptional activity in cells that possessed Y537S ERα^5^. In the WT/D538G heterozygous cells RU39411, fulvestrant, and OP-1074 showed the best IC_50_ values at 0.26 + 0.53, 0.57 ± 0.69, and 0.52 ± 0.63 nM respectively. At the maximum dose: OP1154, PIP, fulvestrant, and OP1074 showed the greatest transcriptional inhibition at normalized fluorescence of 0.25 ± 0.03, 0.29 ± 0.07, 0.32 ± 0.03, and 0.37 ± 0.05 respectively. Only these four molecules returned the D538G transcriptional activity back to their respective WT values. Together, these data suggest that neither the potency nor the efficacy of induced ERα degradation correlate with improved transcriptional inhibition for the Y537S and D538G mutant receptors in this assay.

**Figure 5:**
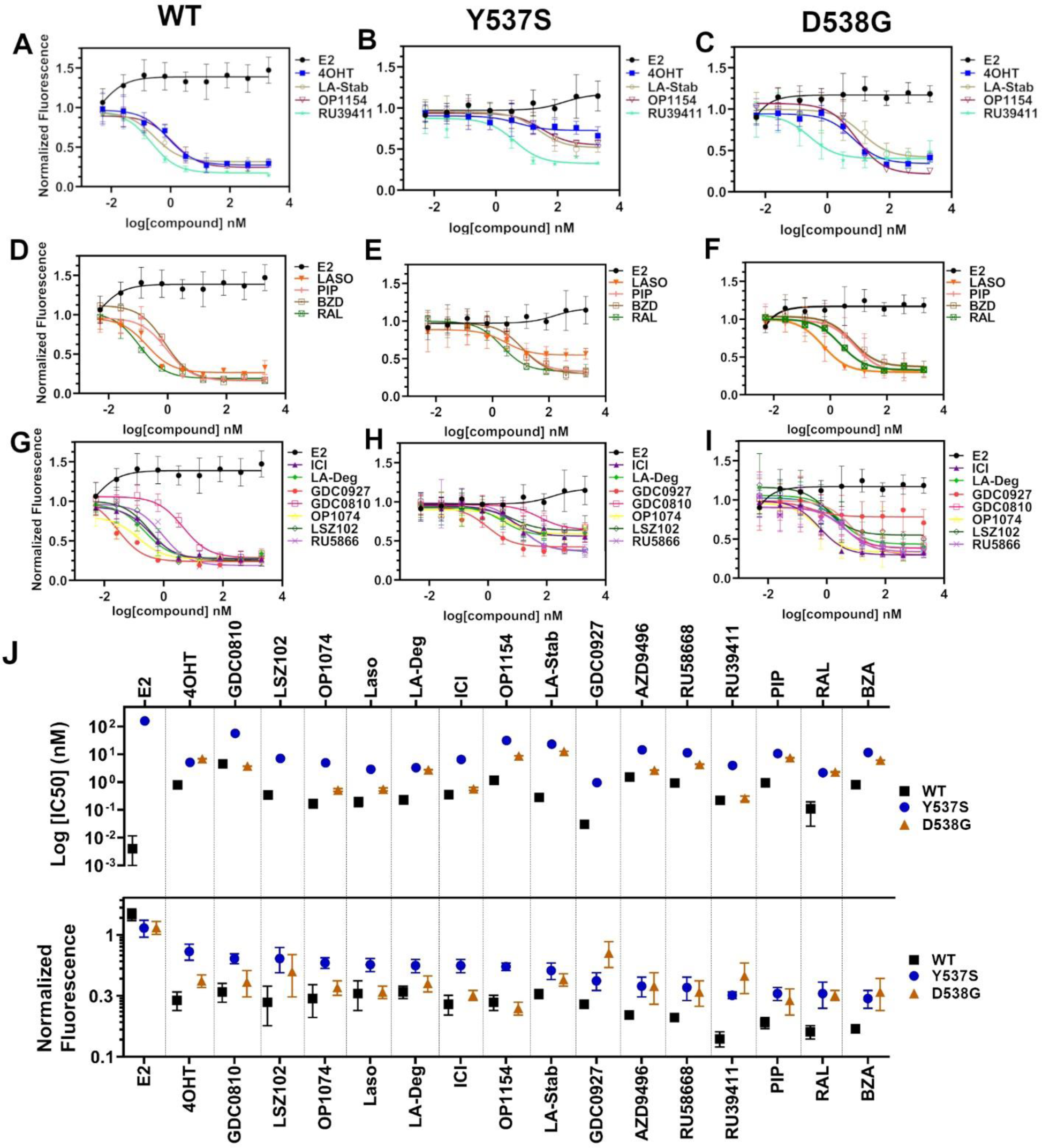
Transcriptional reporter gene assays in MCF7 cells with WT, WT/Y537S, and WT/538G *ESR1*. A) SERMs in WT MCF7 cells. B) SERMs in WT/Y537S MCF7 cells. C) SERMs in WT/D538G MCF7 cells. D) SERM/SERDs in WT MCF7 cells. E) SERM/SERDs in WT/Y537S MCF7 cells. F) SERM/SERDs in WT/D538G MCF7 cells. G) SERDs in WT MCF7 cells. H) SERDs in WT/Y537S MCF7 cells. I) SERDs in WT/D538G MCF7 cells. J) Summary of IC_50_ and normalized fluorescence at 5 μM compound after 24 hours for each compound. Data are ordered from left to right based on normalized fluorescence at highest dose in Y573S MCF7. Poor IC_50_ fits were omitted. Data are shown as the mean of two biologic replicates± standard deviation. All data are normalized to cell count in their respective wells.

### LA-Deg and LA-Stab Show Anti-Proliferative Activities in MCF7 Cells with WT/Y537S *ESR1*

We used label-free cell counting to determine whether induction of ERα degradation improved anti-proliferative outcomes in MCF7 cells with heterozygous WT/Y537S *ESR1.* This model was chosen because it shows the greatest resistance to inhibition by antiestrogens^5^. Cells were treated with 4OHT, fulvestrant, Laso, LA-Stab, or LA-Deg at 1 nM, 50 nM, and 1 μM in the presence of 1 nM E2 alongside vehicle (DMSO) and 1 nM E2-only controls. Subsequently, cells were counted every six hours for 116 hours, with the exception of a gap of 24-hours between hours 84 and 108 when the media was renewed and cells re-treated. The experiment was halted when cells in the E2-treated well reached 100% confluence. All treatments were performed with three biological replicates and a total of 9 technical replicates per ligand per concentration. **Figure 6** shows the proliferation data and a summary of results. As expected, addition of 1 nM E2 enhanced cell proliferation. No compound completely abrogated proliferation, even at the highest dose. Every compound shows similar efficacies at the maximum 1 μM dose, although LA-Deg and LA-Stab appear slightly improved and closest to veh levels, but differences were not statistically significant compared to the other molecules at the same dose. Fulvestrant and Laso on the other hand show improved efficacies at the 1 nM and 50 nM doses compared to the other molecules. Together, these data show that induction of ERα degradation does not by itself predict the anti-proliferative activities of an antiestrogen in this cell model.

**Figure 6:**
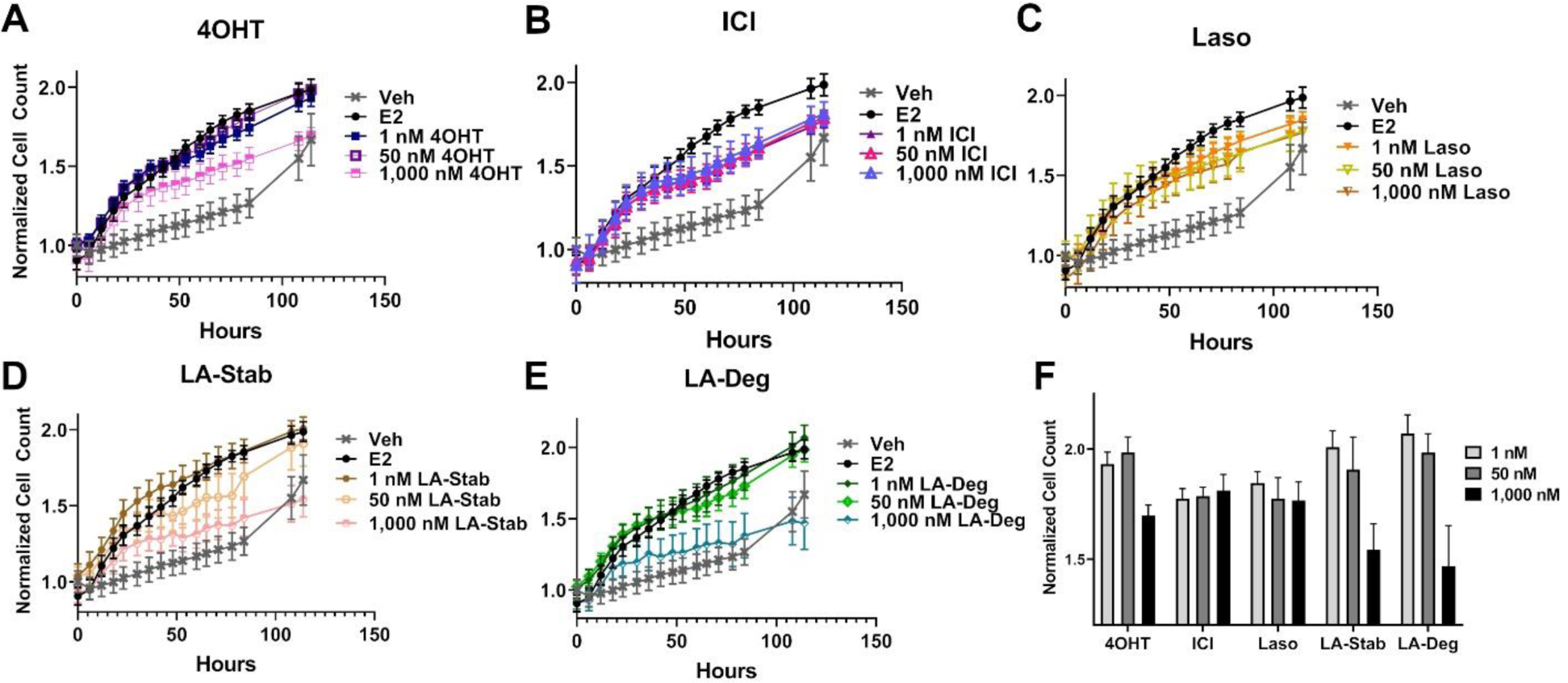
Anti-proliferative activities of SERMs and SERDs in MCF7 cells with heterozygous WT/Y537S *ESR1*. A) 4-hydroxytamoxifen (4OHT). B) Fulvestrant (ICI). C) Lasofoxifene (Laso). D) Lasofoxifene-Stabilizer (LA-Stab). E) Lasofoxifene-Degrader (LA-Stab). F) Normalized cell count after 116 hours. All antagonist treatments are in the presence of 1 nM estradiol (E2). All data are normalized to initial cell count of the vehicle wells in their respective plates. All data are mean ± s.e.m. for three biological replicates and a total of nine technical replicates.

### Structural Basis of Improved Antagonism for Y537S ERα

We solved x-ray crystal structures of antiestrogens in complex with Y537S and WT ERα ligand binding domain (LBD) to understand the structural basis of differential transcriptional antagonistic potencies. We solved x-ray crystal structures for BZA, RAL, 4OHT, LA-Deg, LA-Stab, RU39411 and LSZ102 in complex with the Y537S mutant. We also solved structures with the WT LBD for those molecules with no existing structures (LA-Deg, LA-Stab, RU39411) or were able to solve to improved resolution (RAL). We were unable to solve x-ray crystal structures for antagonists in complex with the D538G mutant, despite earlier success with 4-hydroxytamoxifen^11^. Overall, these structures present canonical ERα LBD head-to-head homodimers with 2 to 4 monomers in the asymmetric unit. As with other ERα LBD structures, these show significant crystal contact influences at H12 of one monomer but not the other. For analysis purposes, we focused on the monomers with the fewest crystal contacts near H12. In the WT LBD: RAL, BZA and LSZ102 showed the fewest H12 contacts at Chain A. For LA-Deg, LA-Stab, and 4OHT it was Chain B and Chain D for RU39411. In the Y537S structures, it was chain A for RAL, BZA, and LSZ102, chain B for LA-Stab and 4OHT, and Chain D for LA-Deg and RU39411.

#### Enforcing WT Antagonist Conformation Enables Efficacious Transcriptional Inhibition of Y537S ERα

RAL, BZA, and RU39411 showed the greatest efficacy of transcriptional inhibition at the maximal doses in MCF7 cells with WT/Y537S *ESR1*. Superposition of WT and Y537S RAL structures, based on alpha carbon positions, shows that H12 maintains a similar binding orientation but appears slightly more buried in the AF2 cleft in the Y537S vs WT (**Figure 7**). The RAL ligand itself shifts towards the loop connecting H11 to H12 (H11-12 loop) and disorders it relative to WT (**Figure 7A**). BZA induces a similar H12 conformation by increasing H12 burial within the AF2 cleft in the Y537S vs the WT (**Figure 7B**). Interestingly, BZA reorients within the Y537S ligand binding pocket to accommodate this new conformation, with its azepan ring shifting towards the H11-12 loop and phenolic ring on its core rotating approximately 60°. This increased burial of the Y537S H12 in the RAL and BZA structures may explain the reduced degrader activities observed for these molecules with this mutant (**Figure 2J**). Likewise, the RU39411 structures show similar H12 conformational changes with H12 moving closer to the AF2 cleft in the Y537S vs WT (**Figure 7C**). We previously observed similar WT and Y537S structures for lasofoxifene, which also maintains significant transcriptional inhibitory activities^28^. For LA-Deg, its 3R-methylpyrrolidine points towards the H11-12 loop and H12 is less ordered, similar to other SERDs^26^. In contrast, the 2S-methylpyrroldine of LA-Stab points into a hydrophobic pocket between H3 and H5 and appears to enhance H12 stability. In the Y537S complexes, H12 of LA-Deg and LA-Stab both shift H12 slightly away from the AF2 cleft (**Figure 7D and E**). LSZ102 and 4OHT showed the greatest reduction in transcriptional inhibition in MCF7 cells with the Y537S mutation. In these structures, H12 is less helical and moves in new solvent-facing vectors away from H3. Additionally, the dimethylamine of 4OHT no longer forms a hydrogen bond with D351 in the Y537S structure (**Figure 7G**).

**Figure 7:**
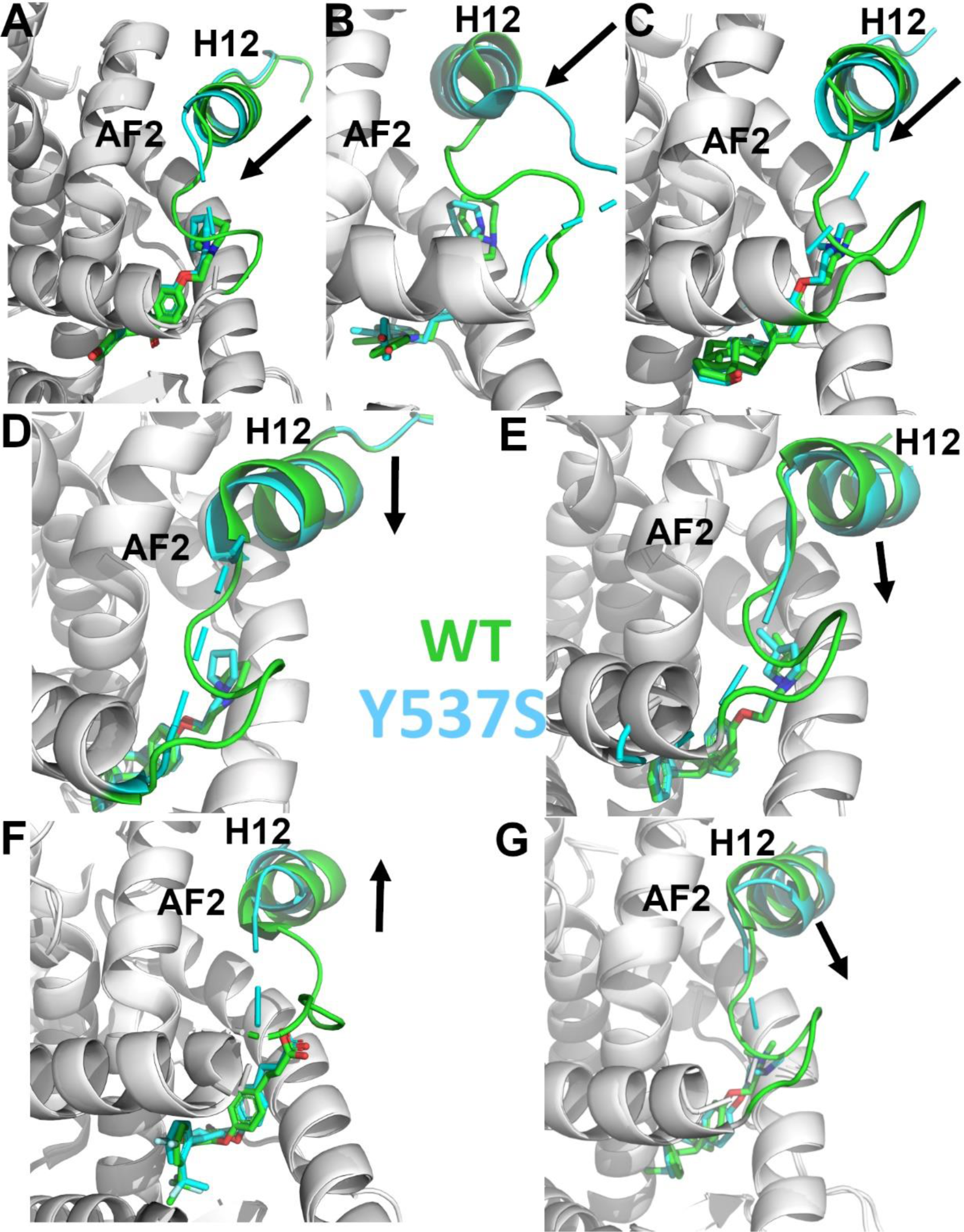
Enforcing helix 12 antagonistic conformation enables potent Y537S ERα transcriptional inhibition. In each panel, arrows show the direction of H12 relocation in the Y537 relative to WT. Helix 11-12 loop, helix 12 and the respective ligands are colored green for WT and cyan for Y537S. All overlays are based on alpha carbon positions. Panels are ordered from most transcriptionally potent (RAL) to least (4OHT) in WT/Y537S *ESR1* MCF7 cells. Protein is shown as cartoon ribbons and ligands as sticks. A) Raloxifene (PDBs: 7KBS and 7KCA). B) Bazedoxifene (PDBs: 4XI3 and 6PSJ). C) RU39411 (PDBs: 6VIG and 6VJ1). D) LA-Stab (PDBs: 6VNN and 6VPK). E) LA-Deg (PDBs: 6VMU and 6VVP). F) LSZ102 (PDBs: 6B0F and 6V8T). G) 4OHT (PDBs: 5W9C and 6V87).

#### Reduced Efficacy for Y537S Correlates with Decreased H12 Surface Area Burial in the AF-2 Cleft

Buried surface area (BSA) calculations for the residues comprising H12 (537-548) in the representative Y537S LBD monomers show that H12 surface area burial within the AF-2 cleft correlates with transcriptional antagonistic efficacies. Proteins, Interfaces, Structures, and Assemblies^35^ was used to calculate BSA where BZA, RAL, and RU39411 show the greatest burial at 379.75, 377.87, and 325.21 Å^2^ respectively. Molecules with poorer potencies showed markedly reduced BSA at 245.57, 56.13, 27.62, and 14.69 Å^2^ for LA-Stab, LSZ-102, LA-Deg, and 4OHT respectively. Together, these data suggest that antiestrogens that enforce a highly buried H12 antagonist conformation in the AF-2 cleft possess the greatest transcriptional repressive activities in Y537S *ESR1* breast cancer cells.

## DISCUSSION

Chemical manipulation of ERα H12 conformation and mobility tunes receptor lifetime in living cells. SERMs extend ERα lifetime, increase receptor accumulation, and are antagonists in the breast and agonists in the bone and uterine epithelium^17^. Tissue-specific partial agonist activities stem from the SERMs inhibition of Activating Function-2 (AF-2) but not AF-1 genomic activities^36^. SERDs decrease the quantity and lifetime of ERα in the cell and are pure antagonists by effectively prohibiting ERα transcriptional activities. Activating somatic mutations near ERα helix 12, Y537S and D538G, enable hormone therapy resistance in metastatic ER+ breast cancers^5, 6^. Antiestrogens with improved potencies including bazedoxifene, fulvestrant, OP-1074 and GDC-0945 possess variable degrees of ERα degrading/downregulating activities but show significant potencies and efficacies in mutant *ESR1* breast cancer cells^5, 25, 26^. Initial studies suggested that SERD activity was necessary for therapeutic potency^5^; however, Lainé *et al.* also showed that the SERM lasofoxifene induced marked anti-cancer activities in xenografts of Y537S *ESR1* breast cancer cells without fulvestrant-level ERα degradation^28^. In addition, Wardell *et al.* showed that antagonism rather than ERα degradation/downregulation drives fulvestrant efficacy^31^. This is supported by observation in cell lines that fulvestrant retains transcriptional inhibition properties even when no increased ERα degradation is observed^37^. Based on this work, we hypothesized that SERD activity by itself may not predict the potency and efficacy of transcriptional suppression in breast cancer cells harboring Y537S or D538G *ESR1*. To understand whether chemical manipulation of ERα cellular lifetime correlates with improved antagonistic activities, we synthesized methylated derivatives of lasofoxifene that uniquely influence receptor cellular lifetime and accumulation. We compared these molecules to a comprehensive panel of SERMs, SERM/SERDs and SERDs for their effects on WT and mutant ERα cellular stability and transcriptional activity in breast cancer cells.

To quantify the influence of antiestrogens on ERα cellular lifetime, we developed a high-throughput system to observe WT, Y537S, or D538G ERα expression in living T47D breast cancer cells. We also quantified differences in ERα SUMOylation, a post-translational modification that is induced by fulvestrant together with ubiquitination and that influences interaction with DNA and transcriptional activity in ER+ MCF7 cells^15^. We discovered that addition of a 3R-methyl group on lasofoxifene’s pyrrolidine induced both ERα SUMOylation and degrading activities, while addition of a 2S-methyl on the pyrrolidine leads to an accumulation of ERα in the breast cancer cells.

This approach also revealed an unexpected effect of each mutant on ERα expression and cellular lifetime. Most SERD molecules exhibited diminished capacities to affect Y537S ERα levels compared to WT. Two carboxylic acid-containing SERD molecules AZD9496 and LSZ102 unexpectedly switched to stabilizing Y537S, similar to 4-hydroxytamoxifen (4OHT). Of note, AZD9496 did not induce SUMOylation with WT ERα, in keeping with a different conformation of the receptor observed in carboxylic acid-containing antiestrogens. Apart from this discrepancy, the general good degree of correlation between the impact of antiestrogens on SUMOylation of WT or mutant ER and their impact on receptor lifetime suggests either a cross-talk between SUMOylation and ubiquitination, possibly mediated by SUMO targeted ubiquitin ligases (STUbLs) such as RNF4, or similar structural determinants for addition of either marks by their respective enzymatic complexes. It will therefore be of interest in the future to explore whether carboxylic acid-containing antiestrogens selectively induce increased receptor ubiquitination, or lead to degradation by other mechanisms.

We were surprised that the Y537S mutant showed nearly identical unliganded/apo and E2-treated profiles, with enhanced cellular expression in the presence hormone to similar levels to WT in the absence of hormone. In addition to constitutive transcriptional activities and resistance to inhibition by tamoxifen, the Y537S also likely contributes to breast cancer pathology through increasing the quantity of active receptor in the cell. The D538G mutant was also surprising because it dampened the influence of SERDs on ERα SUMOylation, accumulation, and lifetime. It suppressed the stabilizing effects of SERMs on WT ERα, correlating with induction of weak SUMOylation of the D538G mutant but not WT receptor. The D538G *ESR1* mutant occurs with the greatest frequency in patients^8^ but its selective advantage has remained unclear. Our studies suggest that the diminished ability of cells to degrade D538G ERα coupled with its low-level constitutive transcriptional activities contributes a pathological advantage within the tumor in the presence of first-line endocrine therapies. Further studies will reveal whether these alterations to ERα lifetime influence breast cancer pathological endpoints and therapeutic response.

Transcriptional reporter gene assays in MCF7 cells harboring WT, WT/Y537S, or WT/D538G *ESR1* were used to understand whether the degrees of ERα degradation and/or SUMOylation correlated with transcriptional inhibitory activities. We identified antiestrogens across the SERM-SERD spectrum that demonstrated significantly improved transcriptional repressive activity over 4OHT in these cells. Interestingly, RU39411, which induced a significant enrichment of ERα levels (SERM-like), showed enhanced transcriptional inhibition compared to most molecules. Despite their unique influences on WT ERα cellular stability, the methylated lasofoxifene derivatives showed identical transcriptional antagonistic efficacies in the WT and mutant cell lines compared to unmodified lasofoxifene.

Cellular proliferation assays in MCF7 cells with WT/Y537S ESR1 were used to reveal the role of induced ERα degradation in anti-proliferative activities. We showed that 4OHT, fulvestrant, Laso, LA-Deg, and LA-Stab can reduce E2-stimulated proliferation. Fulvestrant and Laso show improved efficacies at lower concentrations, while LA-Stab and LA-Deg show improved efficacies at the maximum dose. However, no molecule completely repressed the proliferation of these cells. Together, our BRET, transcription, and proliferation data are consistent with our observations that SERMs have similar efficacies at suppressing cofactor recruitment in WT and *ESR1mut* cells and suggest that a molecule’s influence on ERα cellular lifetime may not directly correlate with its transcriptional antagonistic and anti-proliferative activities in cells with the *ESR1* mutants. Rather, engineered chemical reduction of ERα cellular accumulation is critical for tuning tissue- and gene-specific agonistic/partial agonistic activities^17^ and preventing AF-1 dependent mechanisms of endocrine resistance^38^.

X-ray crystal structures of WT and Y537S ERα LBD in complex with SERMs, SERM/SERDs, and a SERD showed that the most efficacious molecules enforced H12 burial within the AF-2 cleft. We previously showed that BZA resisted the impact of the Y537S mutation on its antagonist H12 conformation, which was perturbed for the less effective 4-hydroxytamoxifen^13^. Of the structures, RU39411 (SERM), RAL (SERM/SERD), and BZA (SERM/SERD) demonstrated the greatest transcriptional inhibitory efficacies in WT/Y537S *ESR1* MCF7 cells. These molecules enforced an equivalent antagonistic H12 conformation that imposed significant surface area burial within the AF-2 cleft in both WT and Y537 ERα LBD. Conversely, molecules with diminished Y537S transcriptional potencies poorly enforced this conformation as H12 adopted vectors away from the AF-2 cleft with reduced H12 burial. These structures show that the most potent molecules for the ERα Y537S maintain significant H12 surface area burial within the AF-2 cleft regardless of ability to induce ERα degradation.

Overall, our studies show how minor chemical modifications lead to switches in ERα stabilizing or degrading activities onto an antiestrogen scaffold, likely via modulation of ERα post-translational modification. This is consistent with antiestrogens exhibiting a whole spectrum of SERD activities, 4OHT and RU39411 having the most stabilizing influence and fulvestrant and RU58,668 the greatest degrading impact^39^. While these activities may not confer an antagonistic benefit, they are important drivers of AF-1 mediated tissue-specific activities^36^. Importantly, we also showed that enforcing H12 burial in the AF-2 cleft improves transcriptional antagonistic activities on a reporter vector in breast cancer cells harboring WT/Y537S *ESR1*. Our studies, combined with the recent work from the McDonnell Laboratory showing that transcriptional antagonism contributes most of fulvestrant’s therapeutic activities independent of maximal receptor depletion^31^, and the observed significant efficacy of the SERM/SERD lasofoxifene for the Y537S mutant^28^, suggest that future antiestrogen development efforts for treatment of tumors with constitutively active mutations should focus on maximizing transcriptional antagonistic activities.

## MATERIALS AND METHODS

### Cell culture

MCF-7 and T47D cells were purchased directly from ATCC. MCF-7 cells were cultured in DMEM supplemented with 10% FBS and 1% penicillin-streptomycin. T47D cells were cultured in RPMI supplemented with 10% FBS and 1% penicillin-streptomycin.

### Live cell assay of Halo-ER Degradation

T47D cells with stable TetON halo-tagged WT, Y537S, or D538G ERαwere plated in a 6 well dish at a density of 15,000 and 30,000 cells, respectively. These cells were cultured for 48 hours in phenol-free media supplemented with charcoal-stripped FBS in puromycin. Following this incubation, 1 µg/mL doxycycline and 1 µM G618 were added to each well. Cells were then treated with increasing concentrations of our compounds of interest (0.00512, 0.0256, 0.128, 0.64, 3.2, 16, 80, 400, 2000, and 5,000 nM) for 24 hours. After treatment, the cells were imaged using an Incucyte S3. ER degradation was quantified by using the red channel integrated intensity per image normalized to the phase channel confluence area. Assays were performed twice with three technical replicates each.

### Chemical Synthesis

**1-[4-(2-Benzyloxy)ethoxy]phenyl-2-bromo-6-methoxy-3,4-dihydronaphthalene (5)** was prepared starting from 4-bromophenol in four steps according to a literature procedurewith slightly modification^40^.

**Supplemental File 8** shows the synthetic route used to generate the lasofoxifene analogs. A mixture of 4-bromophenol (40.0 g, 231 mmol), ethylene carbonate (38.0 g, 432 mmol) and K_2_CO_3_ (68.0 g, 492 mmol) in DMF (200 mL) was heated for 36 h at 100 °C and then cooled. The reaction mixture was poured into water (500 mL), extracted with ether (3 x 400 mL) and the combined organic layers was washed with 5% NaOH (250 mL), brine and dried over Na_2_SO_4_. The solid was filtered off and the solvent was removed by rotary evaporator under reduced pressure to give 2-(4-bromophenoxy)ethanol (**2**) as colorless solid (49.5 g, 99% yield). ^1^H NMR (CDCl_3_/TMS) δ 7.37 (m, 2H), 6.79 (m, 2H), 4.04 (m, 2H), 3.96 (m, 2H), 2.14 (t, 1H, J = 6.0 Hz); ^13^C NMR (CDCl_3_) δ 157.9, 132.5, 116.5, 113.4, 69.5, 61.5 ppm.

Under argon to the solution of **2** (48.8 g, 225 mmol) in dry DMF (650 mL), NaH (95%, 8.53 g, 338 mmol) was added at 0 °C. After stirred at rt for 30 minutes, tetra-*n*-butylammonium iodide (872 mg, 2.36 mmol) and benzyl chloride (29.9 g, 236 mmol) was added. The reaction mixture was stirred at rt overnight and then quenched with saturated aqueous ammonium chloride (150 mL). The product was extracted with dichloromethane (4 x 100 mL). The combined organic layers were washed with brine and dried over Na_2_SO_4_. The solvent was removed and the residue was purified by silica gel chromatography, eluting with 10% ethyl acetate to give 4-[2-(benzyloxy)ethoxy]bromobenzene (**3**) (57.7 g, 74% yield). ^1^H NMR (CDCl_3_/TMS) δ 7.37-7.28 (m, 7H), 6.81 (m, 2H), 4.63 (s, 2H), 4.11 (m, 2H), 3.82 (m, 2H); ^13^C NMR (CDCl_3_) δ 158.1, 138.1, 132.4, 128.6, 127.9, 116.6, 113.2, 73.6, 68.5, 67.8 ppm.

Under argon a mixture of **3** (28 g, 91.2 mmol) and magnesium turnings (2.32 g, 95.8 mmol) in THF (300 mL) was heated under reflux for 24 h with the addition of several iodine particles to initiate the reaction. The solution was cooled to rt and 6-methoxy-3,4-dihydronaphthalen-1(2H)-one (11.7 g, 66.4 mmol) in THF (100 mL) was added dropwise. After the reaction mixture was stirred at rt for 20 hours, the reaction was quenched with 5% HCl (100 mL). The product was extracted with ether, dried over Na_2_SO_4_. The solvent was removed, the residue was isolated by silica gel chromatography, eluting with 10% ethyl acetate in hexane to give 1-[4-(2-benzyloxy)ethoxy]phenyl-6-methoxy-3,4-dihydronaphthalene (**4**) as light-yellow oil (13.9 g, 54% yield). ^1^H NMR (CDCl_3_/TMS) δ 7.40-7.20 (m, 7H), 7.00-6.90 (m, 3H), 6.76 (d, 1H, *J* = 2.4 Hz), 6.63 (dd, 1H, *J* = 8.6, 2.6 Hz), 5.91 (t, 1H, *J* = 4.4 Hz), 4.65 (s, 2H), 4.18 (m, 2H), 3.85 (m, 2H), 3.79 (s, 3H), 2.81 (t, 2H, *J* = 8.0 Hz), 2.36 (m, 2H); ^13^C NMR (CDCl_3_) δ 158.6, 158.1, 139.0, 138.8, 138.2, 133.7, 129.8, 128.6, 127.92, 127.85, 126.7, 124.6, 114.5, 113.9, 110.8, 73.5, 68.7, 67.6, 55.4, 29.0, 23.6 ppm.

Bromine (2.0 mL, 39.5 mmol) was added slowly to the solution of **4** (13.87 g, 35.9 mmol) in THF (120 mL) at 0 °C. After 5 min, triethylamine (5.5 mL, 39.5 mmol) was added while the mixture was stirred vigorously. The reaction mixture was stirred at rt for 10 min. The reaction mixture was washed with 5% Na2S2O3, extracted with ether, washed with brine and dried over Na_2_SO_4_. The solvent was removed and the residue was isolated by silica gel chromatography, eluting with 10% ethyl acetate in hexane to give 1- [4-(2-benzyloxy)ethoxy]phenyl-2-bromo-6-methoxy-3,4-dihydronaphthalene (**5**) as brown solid (15.6 g, 94% yield). ^1^H NMR (CDCl_3_/TMS) δ 7.40-7.24 (m, 5H), 7.15-7.10 (m, 2H), 7.00-6.94 (m, 2H), 6.70 (d, 1H, *J* = 2.4 Hz), 6.60-6.50 (m, 2H), 4.66 (s, 2H), 4.20 (m, 2H), 3.87 (m, 2H), 3.77 (s, 3H), 2.95 (m, 4H); ^13^C NMR (CDCl_3_) δ 158.8, 158.1, 138.2, 137.6, 136.2, 132.4, 131.0, 129.4, 128.6, 127.92, 127.85, 127.5, 120.7, 114.5, 113.7, 110.0, 73.5, 68.7, 67.5, 55.4, 35.2, 30.1 ppm.

#### 1-[4-(2-Benzyloxy)ethoxy]phenyl-6-methoxy-2-phenyl-3,4-dihydronaphthalene (6)

Under argon a mixture of **4** (3.84 g, 8.25 mmol), phenylboronic acid (1.11 g, 9.1 mmol), Pd(PPh3)4 (286 mg, 0.247 mmol) and potassium carbonate (2.28 g, 16.5 mmol) in dry THF (100 mL) was stirred at reflux for 24 h. After the mixture was cooled down, water (30 mL) was added. The organic layer was separated. The aqueous layer was extracted with ether, washed with brine. The organic layers were combined and dried over Na_2_SO_4_. The solvent was removed, the residue was isolated by silica gel chromatography, eluting with 10% ethyl acetate in hexane to give **6** (3.55 g, 93% yield). ^1^H NMR (CDCl_3_/TMS) δ 7.40-7.20 (m, 6H), 7.15-6.90 (m, 6H), 6.82-6.78 (m, 3H), 6.74 (d, 1H, *J* = 8.8 Hz), 6.60 (dd, 1H, *J* = 8.8, 2.8 Hz), 4.66 (s, 2H), 4.13 (m, 2H), 3.83 (m, 2H), 3.82 (s, 3H), 2.95 (m, 2H), 2.79 (m, 2H); ^13^C NMR (CDCl_3_) δ 158.5, 157.5, 143.4, 138.2, 137.8, 134.9, 134.4, 132.4, 132.2, 130.6, 128.5, 128.4, 127.9, 127.8, 127.7, 127.6, 125.8, 114.2, 113.3, 110.9, 73.5, 68.6, 67.4, 55.4, 30.9, 29.1 ppm.

#### 1-[4-(2-Hydroxyethoxy]phenyl-6-methoxy-2-phenyl-1,2,3,4-tetrahydronaphthalene (7)

To the solution of **6** (3.47 g, 7.5 mmol) in a mixed solvent of ethanol/ethyl acetate (2/1, v/v, 150 mL), palladium hydroxide on carbon (20% wt., 263 mg, 0.375 mmol) was added. After exchange the air with argon, the mixture was then stirred under a hydrogen balloon overnight. TLC showed no starting material left. The catalyst was filtered off. The solvent was removed and the residue was isolated by silica gel chromatography, eluting with 20% ethyl acetate in hexane to give **7** as a white foam (2.105 g, 75% yield). ^1^H NMR (CDCl_3_/TMS) δ 7.15-7.05 (m, 3H), 6.85-6.70 (m, 4H), 6.63 (dd, 1H, *J* = 8.4, 2.8 Hz), 6.49 (d, 2H, *J* = 8.8 Hz), 6.30 (d, 2H, *J* = 8.4 Hz), 4,21 (d, 1H, *J* =6.0 Hz), 3.83 (m, 2H), 3.79 (m, 2H), 3.73 (s, 3H), 3.32 (m, 1H), 3.02 (m, 2H), 2.89 (*brs*, 1H), 2.14 (m, 1H), 1.78 (m, 1H); ^13^C NMR (CDCl_3_) δ 157.7, 156.6, 144.2, 137.6, 134.9, 132.1, 131.4, 131.3, 128.1, 127.6, 125.9, 112.84, 112.82, 112.5, 68.9, 61.1, 55.0, 50.1, 45.3, 30.0, 21.8 ppm. HRMS calcd for C_25_H_30_NO_3_ [MNH_4_^+^] 392.2226, found 392.2222.

#### 1-[4-(2-Mesoxyethoxy]phenyl-6-methoxy-2-phenyl-1,2,3,4-tetrahydronaphthalene (8)

Triethylamine (0.63 mL, 4.52 mmol) was added to a solution of **7** (848 mg, 2.26 mmol) and MsCl (0.35 mL, 4.52 mmol) in THF (25 mL) at 0 °C. The reaction mixture was stirred at rt for 1 h. The reaction was quenched with water, extracted with ethyl acetate, and dried over MgSO_4_. The solvent was removed and the residue was purified by silica gel chromatography, eluting with 20% ethyl acetate in hexane to give the mesylate **8** (0.90 g, 94% yield). ^1^H NMR (CDCl_3_/TMS) δ 7.18-7.10 (m, 3H), 6.85-6.70 (m, 4H), 6.66 (dd, 1H, *J* = 8.4, 2.4 Hz), 6.50 (d, 2H, *J* = 8.4 Hz), 6.33 (d, 2H, *J* = 8.4 Hz), 4,44 (m, 2H), 4.23 (d, 1H, *J* = 4.8 Hz), 4.05 (m, 2H), 3.78 (s, 3H), 3.36 (m, 1H), 3.06 (m, 2H), 2.99 (s, 3H), 2.16 (m, 1H), 1.82 (m, 1H); ^13^C NMR (CDCl_3_) δ 157.9, 156.0, 144.2, 137.8, 135.7, 132.0, 131.50, 131.45, 128.1, 127.8, 126.0, 113.0, 112.9, 112.6, 68.3, 65.6, 55.2, 50.2, 45.3, 37.6, 30.1, 21.9 ppm. HRMS calcd for C_26_H_28_O_5_SNa [MNa^+^] 475.1555, found 475.1550.

#### Typical procedure for the synthesis of 9a-9d

The synthesis of **9a** is representative: To the solution of **8** (85 mg, 0.20 mmol) in anhydrous DMF (5 mL), (3*R*)-3-methylpyrrolidine-HCl salt (242 mg, 2.0 mmol) and *i*-Pr2NEt (700 uL, 4.0 mmol) were added. After the mixture was stirred at 80 °C for 24 h, TLC showed the reaction was completed. The solvent was removed, and the residue was dissolved into ether, washed with saturated NaHCO3, brine, and dried over Na_2_SO_4_. After filtration, the solvent was removed, the residue was dried under vacuum overnight, and then redissolved into dry dichloromethane (10 mL). Under argon the dichloromethane solution was cooled to -78 °C. BBr3 (1.0 M in CH_2_Cl_2_, 2.0 mL, 2.0 mmol) was added. After the reaction mixture was warmed and stirred at 0 °C for 2 h, the reaction was quenched with Na2S2O3 solution at 0 °C. The mixture was basified with 2 N NaOH. extracted with CH_2_Cl_2_, washed with brine and dried over MgSO_4_. The solvent was removed, the residue was purified by silica gel chromatography, eluting with 4% methanol in dichloromethane containing 1% triethylamine to give product **9a** (31 mg, 36% yield). ^1^H NMR (CDCl_3_/TMS) δ 7.15 (m, 3H), 6.80-6.62 (m, 4H), 6.58 (dd, 1H, *J* = 8.4, 2.4 Hz), 6.42 (m, 2H), 6.26 (m, 2H), 4.19 (d, 1H, *J* = 5.2 Hz), 3.98 (m, 2H), 3.35 (m, 1H), 3.15-2.55 (m, 7H), 2.45-2.25 (m, 1H), 2.25-2.05 (m, 3H), 1.76 (m, 1H), 1.42 (m, 1H), 1.05 (d, 3H, *J* = 6.8 Hz); ^13^C NMR (CDCl_3_) δ 156.7, 155.4, 144.6, 137.6, 135.0, 131.5, 131.3, 130.8, 128.30, 127.8, 126.0, 115.15, 115.12, 114.2, 112.8, 65.8, 62.5, 62.3, 55.5, 54.6, 54.4, 50.3, 45.5, 32.43, 32.39, 31.9, 31.8, 30.0, 22.0, 19.9 ppm; HRMS calcd for C_29_H_34_NO_2_ [MH^+^] 428.2590 found 428.2590. HPLC analysis and purification with Chiral pack IBN-3 column, 0.75 mL/min, mobile phase: 0.1% Et_2_NH in MeOH. UV 279 nm, **9a-1**, retention time 8.05 min, **9a-2**, retention time 9.62 min.

**9b** (30 mg, 35% yield) was prepared from **8** (85 mg, 0.20 mmol), (3*S*)-3-methylpyrrolidine-HCl salt (121 mg mg, 1.0 mmol) and *i*-Pr_2_NEt (348 uL, 2.0 mmol) according to the typical procedure: ^1^H NMR (CDCl_3_/TMS) δ 7.14 (m, 3H), 6.80-6.62 (m, 4H), 6.55 (dd, 1H, *J* = 8.4, 2.8 Hz), 6.37 (d, 2H, *J* = 8.8 Hz), 6.26 (d, 2H, *J* = 7.6 Hz), 4.19 (d, 1H, *J* = 4.8 Hz), 3.96 (m, 2H), 3.35 (m, 1H), 3.15-2.75 (m, 6H), 2.60 (m, 1H), 2.40-2.25 (m, 1H), 2.20-2.00 (m, 3H), 1.75 (m, 1H), 1.40 (m, 1H), 1.05 (d, 3H, *J* = 6.8 Hz); ^13^C NMR (CDCl_3_) δ 156.8, 155.5, 144.7, 137.6, 134.9, 131.6, 131.4, 130.7, 128.4, 127.8, 126.0, 115.23, 115.18, 114.3, 112.8, 65.9, 62.6, 62.4, 55.63, 55.60, 54.6, 54.4, 50.3, 45.6, 32.51, 32.45, 31.9, 31.8, 30.1, 22.0, 20.1, 20.0 ppm; HRMS calcd for C29H34NO2 [MH^+^] 428.2590 found 428.2589. HPLC analysis with Chiral pack IBN-3 column, 0.75 mL/min, mobile phase: 0.1% Et_2_NH in EtOH. UV 279 nm, **9b-1**, retention time 5.87 min, **9b-2**, retention time 6.67 min.

**9c** (52 mg, 53% yield) was prepared from **8** (100 mg, 0.23 mmol), (2*R*)-2-methylpyrrolidine-HCl salt (280 mg, 2.3 mmol) and *i*-Pr_2_NEt (800 uL, 4.6 mmol) according to the typical procedure: ^1^H NMR (CDCl_3_/TMS) δ 7.15 (m, 3H), 6.80-6.62 (m, 4H), 6.55 (m, 1H), 6.46 (d, 1H, *J* = 8.4 Hz), 6.41 (d, 1H, *J* = 8.8 Hz), 6.27 (m, 2H), 4.19 (d, 1H, *J* = 4.8 Hz), 3.98 (m, 2H), 3.35-3.15 (m, 3H), 3.02-2.90 (m, 2H), 2.50-2.40 (m, 2H), 2.29 (m,1H), 2.20-2.00 (m, 1H), 2.00-1.90 (m, 1H), 1.90-1.60 (m, 3H), 1.55-1.40 (m, 1H), 1.19 (m, 3H); ^13^C NMR (CDCl_3_) δ 156.8, 155.3, 155.2, 144.63, 144.60, 137.7, 135.0, 134.9, 131.6, 131.5, 131.4, 131.0, 130.8, 128.32, 128.29, 127.8, 126.0, 115.3, 115.2, 114.4, 114.2, 112.9, 112.8, 66.4, 66.2, 61.1, 60.9, 54.5, 54.4, 53.0, 50.3, 45.6, 32.33, 32.26, 30.1, 22.0, 21.8, 21.7, 18.5, 18.4 ppm.; HRMS calcd for C29H34NO2 [MH^+^] 428.2590 found 428.2591. HPLC analysis with Chiral pack IBN-3 column, 0.75 mL/min, mobile phase: 0.1% Et_2_NH in EtOH. UV 279 nm, **9c-1**, retention time 5.88 min, **9c-2**, retention time 6.57 min.

**9d** (59 mg, 60% yield) was prepared from **8** (100 mg, 0.23 mmol), (2*S*)-2-methylpyrrolidine-HCl salt (280 mg, 2.3 mmol) and *i*-Pr_2_NEt (800 uL, 4.6 mmol) according to the typical procedure: ^1^H NMR (CDCl_3_/TMS) δ 7.15 (m, 3H), 6.80-6.62 (m, 4H), 6.55 (m, 1H), 6.47 (d, 1H, *J* = 8.8 Hz), 6.42 (d, 1H, *J* = 8.8 Hz), 6.28 (m, 2H), 4.19 (d, 1H, *J* = 4.8 Hz), 3.98 (m, 2H), 3.40-3.15 (m, 3H), 3.02-2.90 (m, 2H), 2.50-2.40 (m, 2H), 2.26 (m,1H), 2.20-2.15 (m, 1H), 2.00-1.87 (m, 1H), 1.87-1.60 (m, 3H), 1.55-1.40 (m, 1H), 1.16 (m, 3H); ^13^C NMR (CDCl_3_) δ 156.8, 155.31, 155.26, 144.62, 144.60, 137.71, 137.69, 135.0, 134.9, 131.6, 131.5, 131.3, 130.9, 130.8, 128.30, 128.28, 127.8, 126.0, 115.3, 115.2, 114.4, 114.3, 112.9, 112.8, 66.5, 66.3, 61.0, 60.8, 54.6, 54.4, 53.0, 50.3, 45.6, 32.34, 32.28, 30.1, 22.0, 21.8, 21.7, 18.6, 18.4ppm; HRMS calcd for C29H34NO2 [MH^+^] 428.2590 found 428.2591. HPLC analysis and purification with Chiral pack IBN-3 column, 0.75 mL/min, mobile phase: 0.1% Et_2_NH in MeOH. UV 279 nm, **9d-1**, retention time 7.94 min, **9d-2**, retention time 9.11 min.

#### BRET assays

HEK293 HTS cells were maintained at 37℃ in Dulbecco’s modified Eagle’s medium (DMEM) (Wisent) supplemented with 10% fetal bovine serum (FBS) (Sigma), 2% L-glutamine (Wisent) and 1% penicillin-streptomycin (Wisent). Two days prior to transfection, HEK293 HTS cells were washed twice in PBS and seeded in 15 cm petri dishes at a density of 5 x 10^6^ in phenol red-free DMEM supplemented with 1charcoal-stripped FBS (Sigma) and 1% penicillin-streptomycin. Forty-eight hours later, cells were co-transfected with expression plasmids for WT, Y537S, or D538G ERα fused to RlucII and YFP-SUMO3^16, 33^ or SRC1/NCOA1 RID-Topaz YFP^34^. Transfections were performed using linear polyethylenimine (Polysciences Inc.) at a ratio of 3 μg of PEI to 1 μg of DNA per 1.25 x 10^6^ cells and aliquoted at 125 000 cells per well in white 96 well assay plates (Corning). Bioluminescence resonance energy transfer (BRET) assays were carried out 48 h post-transfection by replacing cell media by 1 x HBSS (Wisent) supplemented with 4.5 g/L dextrose and either compounds or vehicle and incubating for 2-3 h at 37℃. For dose-response experiments, compounds were serially diluted 1:3 in 1 x HBSS from a maximum concentration of 9 μM.

Coelenterazine H (Nanolight Technologies) was added to a final concentration of 5 μM and readings were taken following a 5 min incubation at room temperature using a Mithras LB 940 microplate reader (Berthold Technologies). Net BRET ratios were calculated as described previously^16^. All experiments were performed with three to five biological replicates comprised of three technical replicates each. Best-fit EC50 and BRETMax values were determined with Prism. Curves represent average values +/-SEM.

#### Live cell assay of ERE Transcriptional Response

MCF-7 cells with WT, WT/Y537S, and WT/D538G with a 3x-ERE-GFP reporter gene construct with a CMV promoter were plated in a 6 well dish at a density of 15,000 and 30,000 cells, respectively. Cells were cultured for 48 hours in media supplemented with charcoal-stripped FBS. Subsequently, cells were treated with increasing concentrations of compounds of interest (0.00512, 0.0256, 0.128, 0.64, 3.2, 16, 80, 400, 2000, and 5,000 nM) for 48 hours. After treatment, the cells were imaged using an Incucyte S3. ERE transcriptional response was quantified by using the green channel integrated intensity per image normalized to the phase channel confluence area. Assays were performed twice with three technical replicates each.

#### Cellular Proliferation Assay

A BioTek Cytation 5 with BioSpa was used for automatic cell counting experiments. MCF7 WT/Y537S cells were plated in 96-well plates at 750 cells per well in phenol-free medium. After 24 hours, cells placed in charcoal-stripped FBS and allowed to acclimate for 48 hours. Subsequently, cells were treated with 1, 50, and 1,000 nM antiestrogen and 1 nM E2 then placed in the BioSpa. Cells were automatically imaged by bright-field and phase contrast every 6 hours for a total of 116 hours. Cell counts were analyzed using the label-free cell counting protocol on BioTek Gen5 software. These assays were performed three independent times with three technical replicates each.

#### Protein Expression and Purification

A gene containing a hexa-His-TEV fusion of the ERα LBD, residues 300-550 with C381S, C417S, C530S, and L536S in pET21(a)+ was used for all WT ERα LBD x-ray crystal structures, as this construct enables the adoption of a canonical antagonist conformation of the receptor^26^. An identical construct with but with an intact 536L and Y537S was generated using Q5 site-directed mutagenesis (Promega) and was used for all Y537S x-ray crystal structure determinations. *E.coli* BL21(DE3) were used for all recombinant protein expression. An overnight culture at 37°C was inoculated with a single colony in 50 mL LB broth containing 100 μg/mL ampicillin. 5 mL of the overnight culture was used to inoculated each of 10 L of LB-ampicillin, which were allowed to grow at 37°C with shaking until an OD600 = 0.6 was reached, indicating log-phase growth. Protein expression commenced upon the addition of 0.3 mM IPTG and continued overnight at 16°C. Cells were harvested by centrifugation at 4,000 *g* for 15 minutes. Cells were resuspended in 5 mL/g cell paste in 50 mM HEPES pH 8.0, 250 mM NaCl, 20 mM imidazole pH 8.0, 0.5 mM TCEP, and 5% glycerol with the addition of 1 EDTA-free cOmplete protease inhibitor cocktail tablet per 50 mL lysate (Roche). Cells were lysed by sonication on ice with stirring and the lysates were clarified by centrifugation at 20,000 *g* for 30 minutes at 4°C. The clarified lysate was loaded onto 2.5 mL of pre-equilibrated Ni-NTA resin and washed with 5 column volumes of 50 mM HEPES pH 8.0, 250 mM NaCl, 20 mM imidazole pH 8.0, 0.5 mM TCEP, and 5% glycerol. Protein was eluted with 50 mM HEPES pH 8.0, 250 mM NaCl, 400 mM imidazole pH 8.0, 0.5 mM TCEP, and 5% glycerol. A 1:200 mol:mol ratio of hexa-His-TEV protease was added to the eluent and the solution was dialyzed in 4 L of 50 mM HEPES pH 8.0, 250 mM NaCl, 20 mM imidazole pH 8.0, 0.5 mM TCEP, and 5% glycerol overnight at 4° with stirring. ERα was separated from the hexa-His tag and His-TEV protease by passing it over pre-equilibrated Ni-NTA resin and collecting the flow-through. The protein was concentrated to 5 mL, but no higher than 15 mg/mL and placed over a Superdex 200 HiLoad 200 16/600 size exclusion column that was equilibrated with 25 mM HEPES pH 8.0, 150 mM NaCl, 0.5 mM TCEP, and 5% glycerol. A small precipitate peak was observed at the leading edge of the column followed by a well-resolved single peak corresponding to the correct molecular weight of approximately 30,000 Da for ERα LBD. Fractions corresponding to these proteins were concentrated to ≥ 10 mg/mL, flash frozen, and stored at -80°C for later use.

#### X-Ray Crystal Structure Determination

Purified LBD was incubated with 2 mM of each compound between 4 and 16 hours at 4°C prior to crystal screens. The mixture was centrifuged at 20,000 *g* for 30 minutes at 4°C to remove precipitated ligand/protein. Hanging drop vapor diffusion was used to generate all crystals. 2 μL protein at 5 to 15 mg/mL was mixed with 2 μL mother liquor using Hampton VDX plates (Hampton Research, HR3-140). All crystals were grown in 5-20% PEG 3,350 or PEG 8,000, pH 6-8.0, 200 mM MgCl2. For each complex, clear crystals grew at room temperature. Crystals emerged between 16 hours and 2-weeks. For BZA, RAL, 4-OHT, LASO, 48-2, and LSZ102 complexes these crystals appeared as hexagonal pucks. For RU39411 and 49-2 complexes crystals were rhombohedral. Crystals were either directly frozen in mother liquor or incubated in mother liquor with the addition of 25% glycerol as cryo-protectants. All x-ray data sets were collected at the Advanced Photon Source, Argonne National Laboratories, Argonne, Illinois on the SBC 19-BM beamline (0.97 Å). **Figure 7-figure supplement 1** shows representative 2mFo-DFc difference maps for ligands in the ligand binding pocket. **Figure 7-supplemental file 2** contains data collection and refinement statistics.

## ACKNOWLEDGEMENTS

Funding from Susan G. Komen Foundation CCR19608597 (SWF), Ludwig Fund for Metastasis Research (GLG), and Canadian Institutes of Health Research (SM). Results shown in this report are derived from work performed at Argonne National Laboratory (ANL), Structural Biology Center (SBC) at the Advanced Photon Source (APS), under U.S. Department of Energy, Office of Biological and Environmental Research contract DE-AC02-06CH11257.

## SUPPLEMENTAL FILES

**Figure 2-figure supplement 1:**
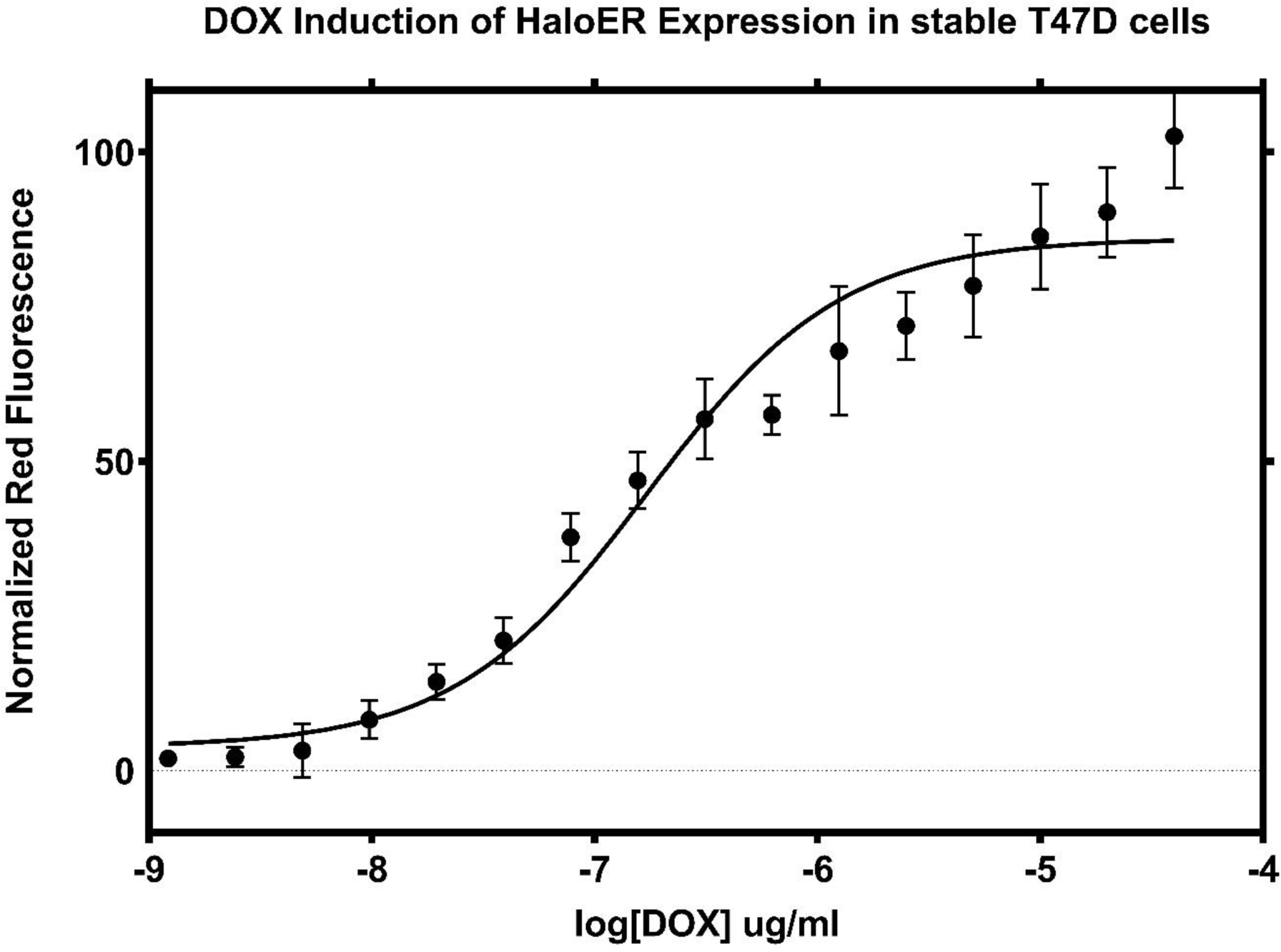
Doxycycline induction of Halo-ERα after 24 hours. Data are representative of the mean of three replicates ± standard error.

**Figure 2-supplemental file 2:**
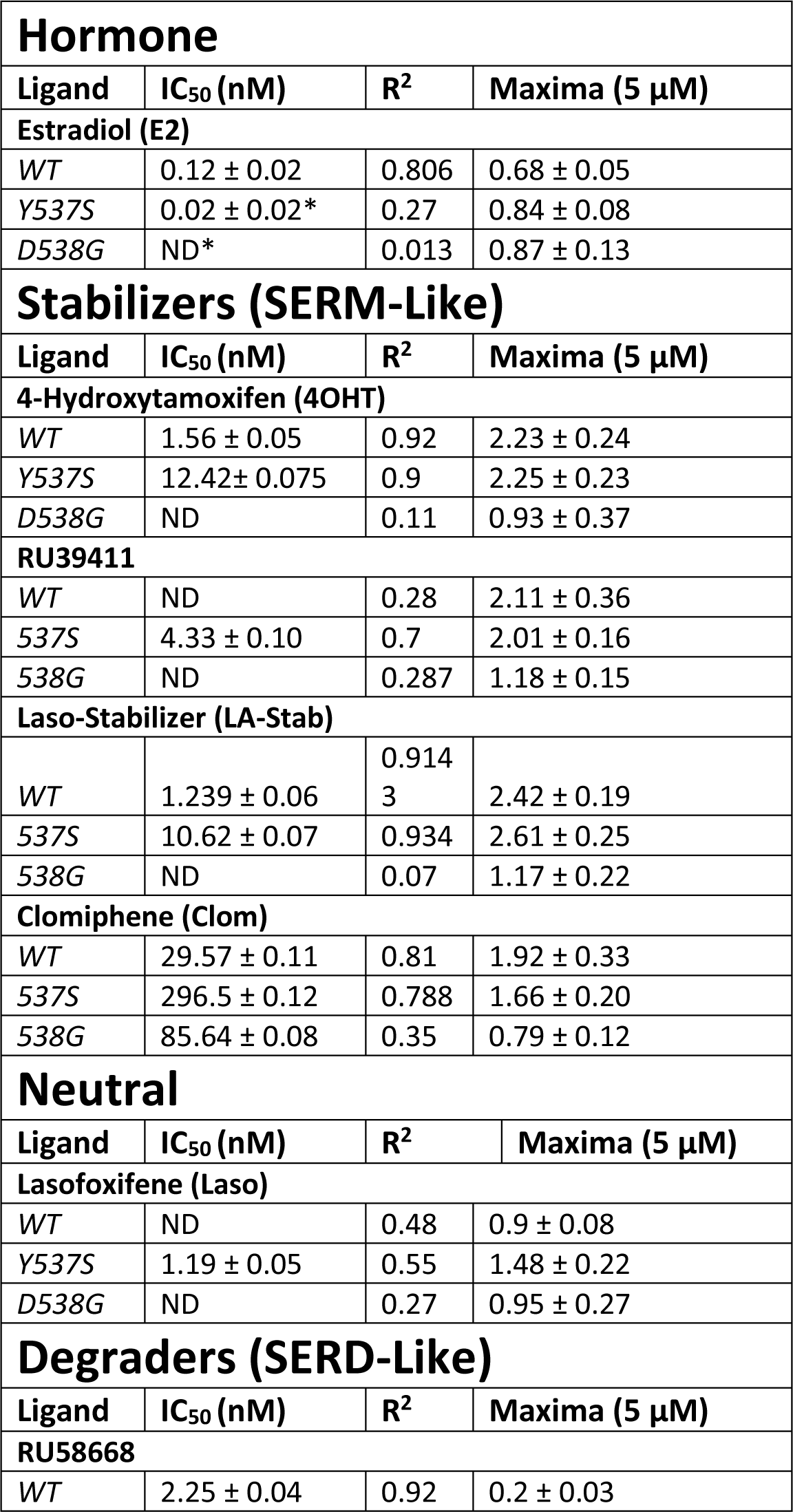
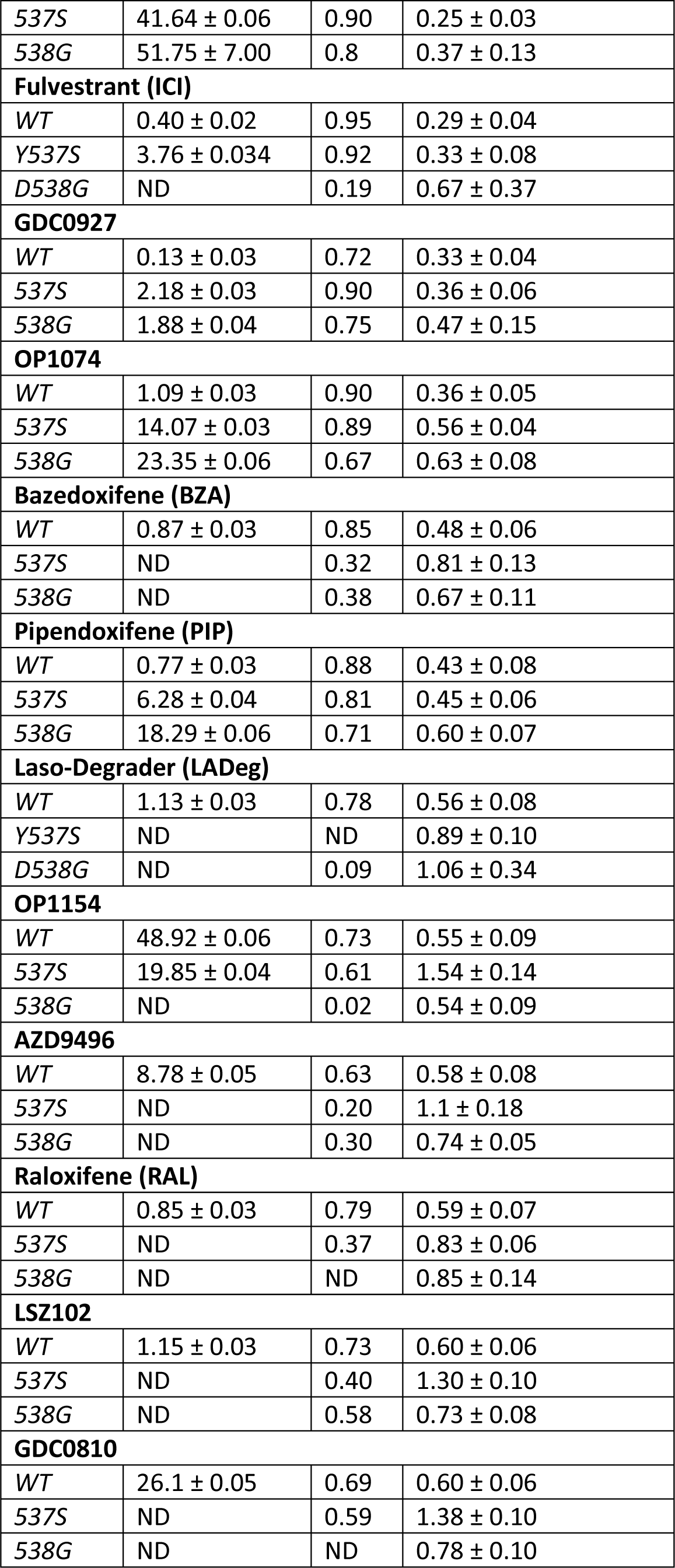
Ligand and Mutational Influences on Estrogen Receptor Alpha Cellular Turnover after 24 Hours. Data were normalized by cell count. Antiestrogens are categorized as stabilizers if their maximum signal for WT ERα at 5 µM is greater than 1.5, neutral if between 0.9 and 1.1, and degrader if less than 0.9 in this assay.

**Figure 2-figure supplement 3:**
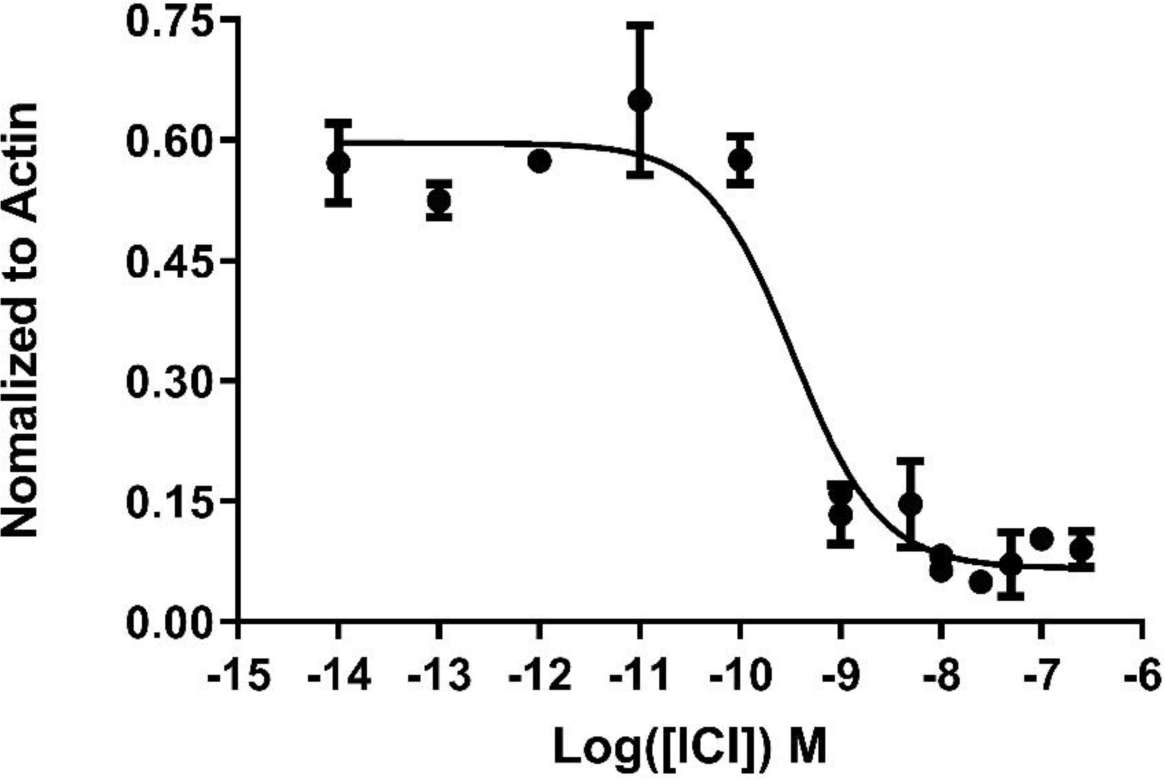
IC50 of fulvestrant (ICI) in normal T47D cells. Data are representative of three replicates per concentration ± standard deviation. All data were normalized to actin control in the individual treatments.

**Figure 3-figure supplement 1:**
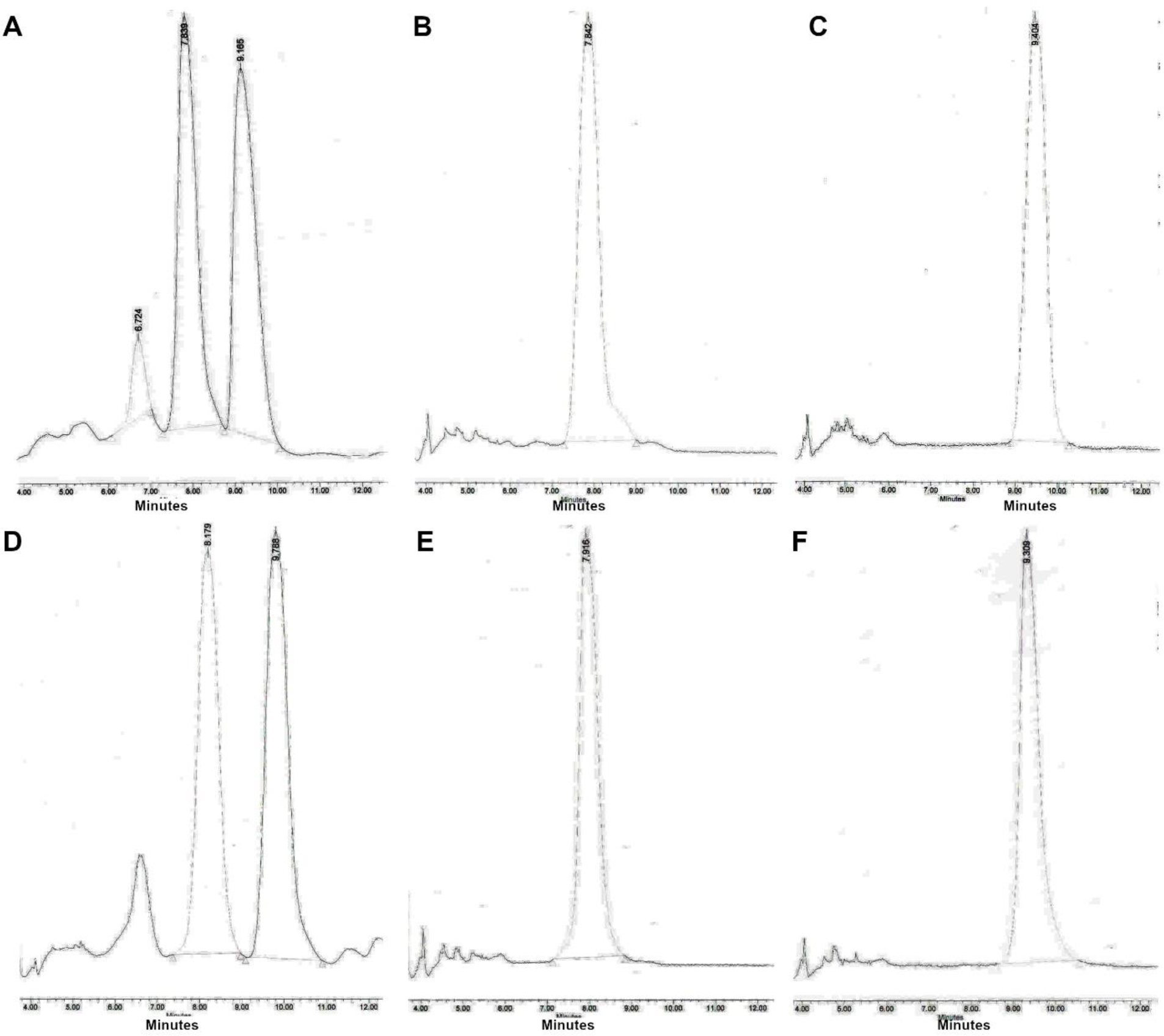
Chiral separation of 3R (LA-Deg) and 2S-methylpyrrolidine (LA-Stab) lasofoxifene derivatives. A) Chromatogram of LA-Deg chiral separation. B) Purification of the LA-Deg peak 1. C) Purification of LA-Deg peak 2. D) Chromatogram of LA-Stab chiral separation. E) Purification of LA-Stab Peak 1. F) Purification of LA-Stab peak 2.

**Figure 4-supplemental file 1:**
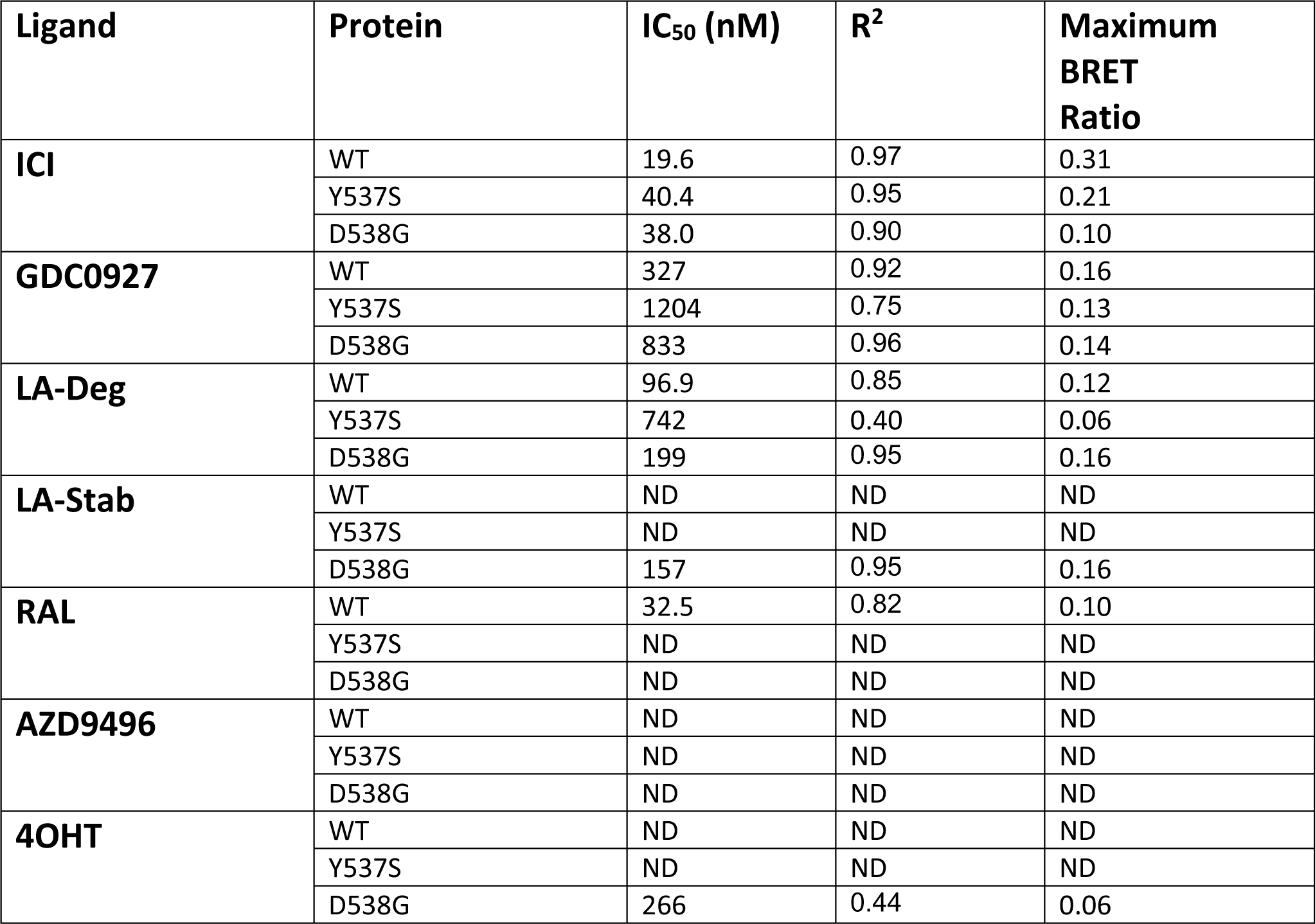
IC50 and Maximum BRET Ratio of Antiestrogen-induced ERα SUMOylation. The s.e.m. values for IC_50_s were all within 20% except for Ral with WT (21.4%) and OHT for D538G (26.2%).

**Figure 4-figure supplement 2:**
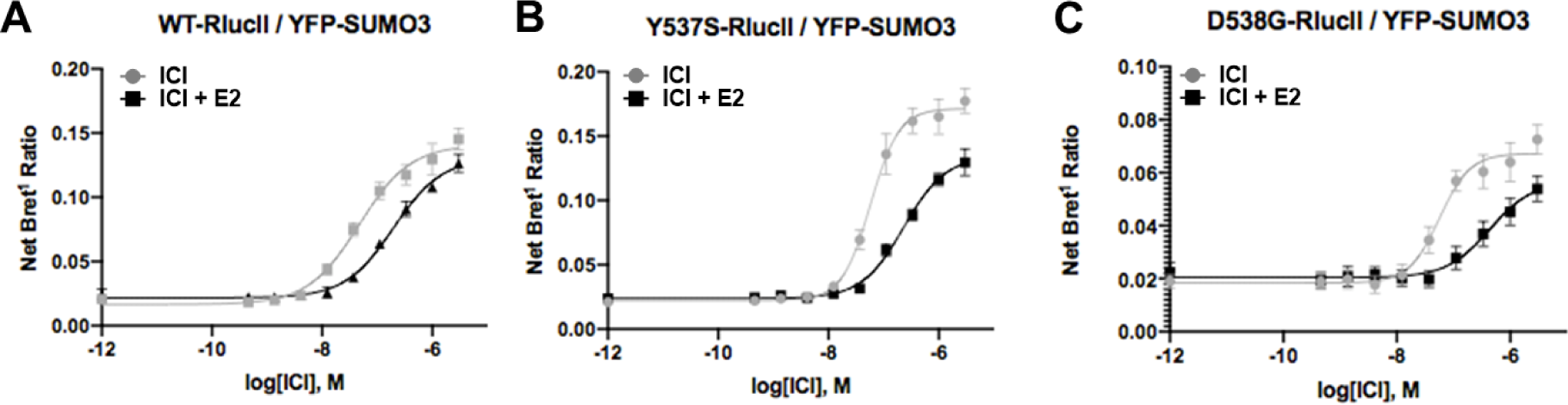
Fulvestrant-induced SUMOylation WT (A), Y537S (B), and D538G (C) ERα in the presence and absence of 5 nM estradiol (E2). Data are mean of three biological replicates ± standard deviation.

**Figure 4-supplemental file 3:**
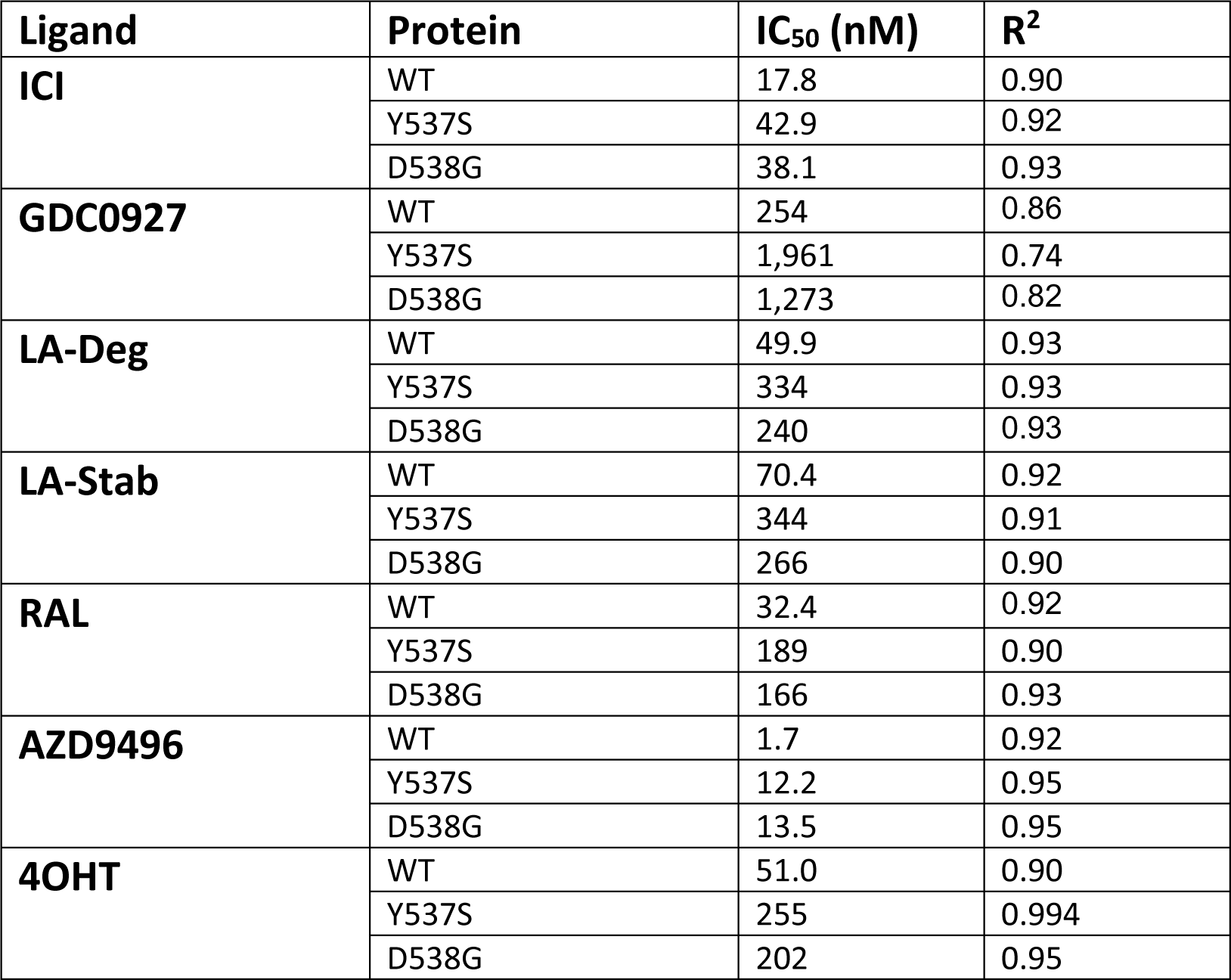
IC50 of SRC1 receptor interacting domain binding to WT and mutant ERα. The s.e.m. values for IC_50_s were all within 50% except for D538G with AZD9496 (52%).

**Figure 5-supplemental file 1:**
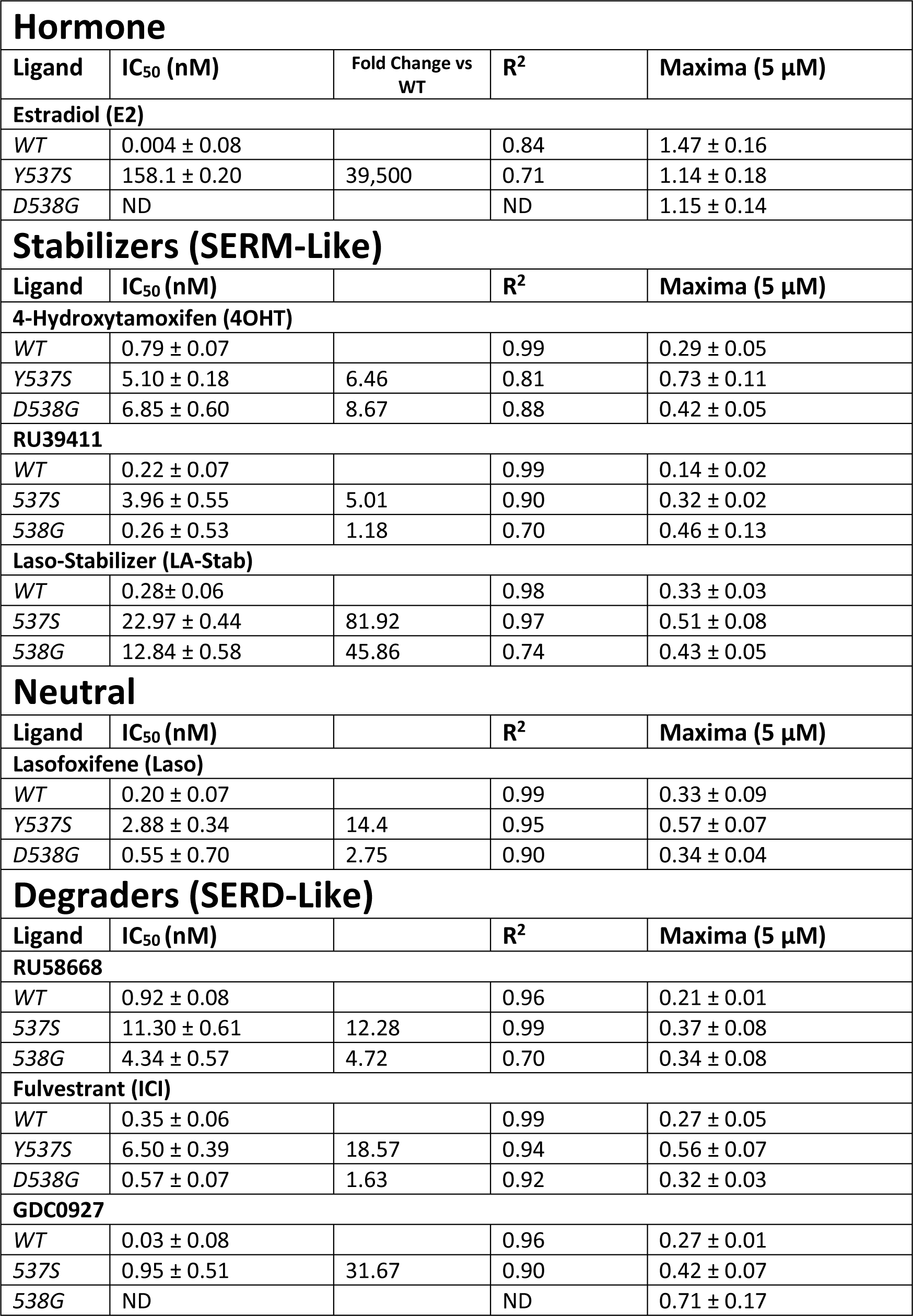
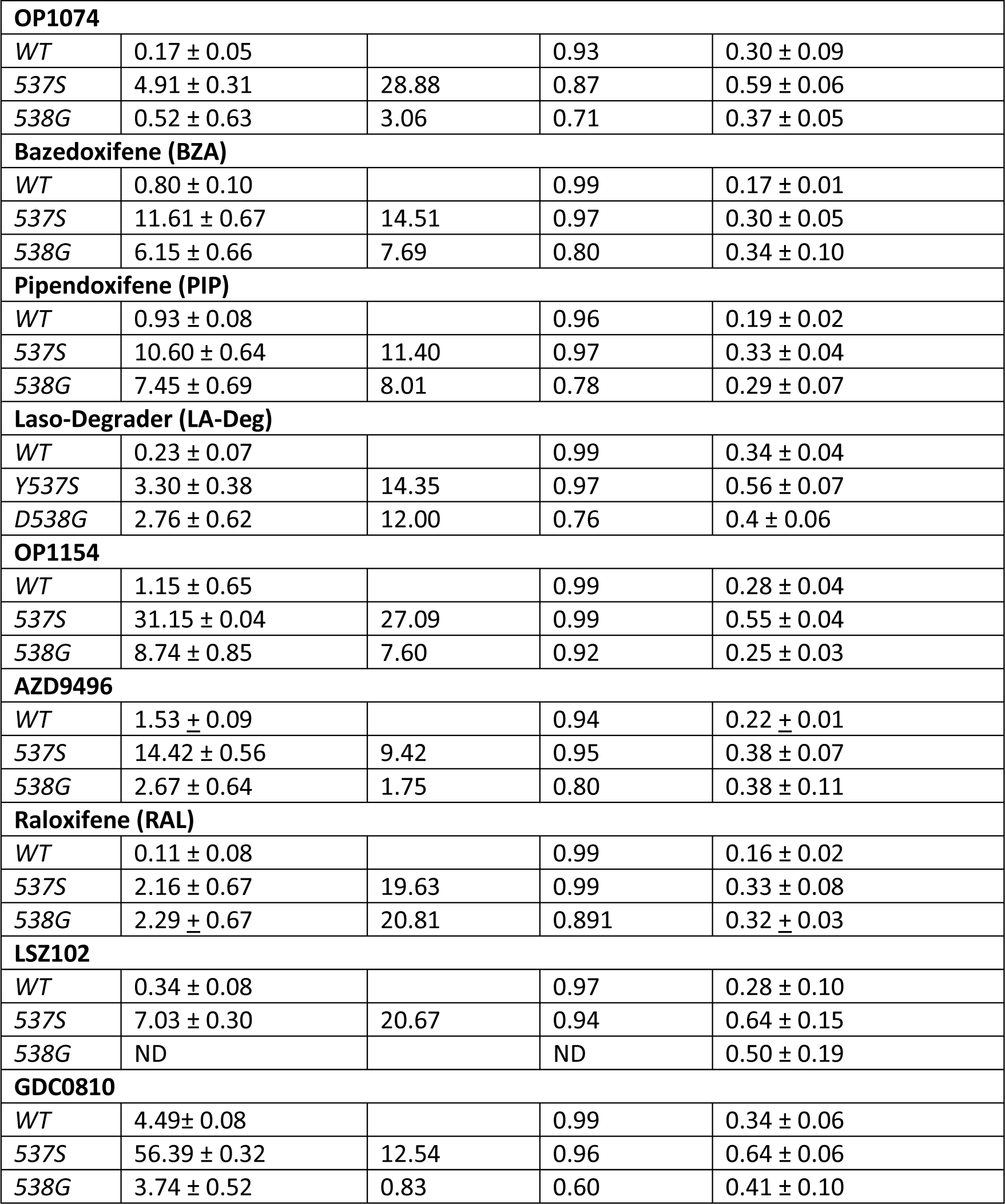
Ligand and Mutational Influences on Estrogen Receptor Alpha Reporter Gene Transcription after 24 Hours. Data were normalized by cell count.

**Supplemental File 8:**
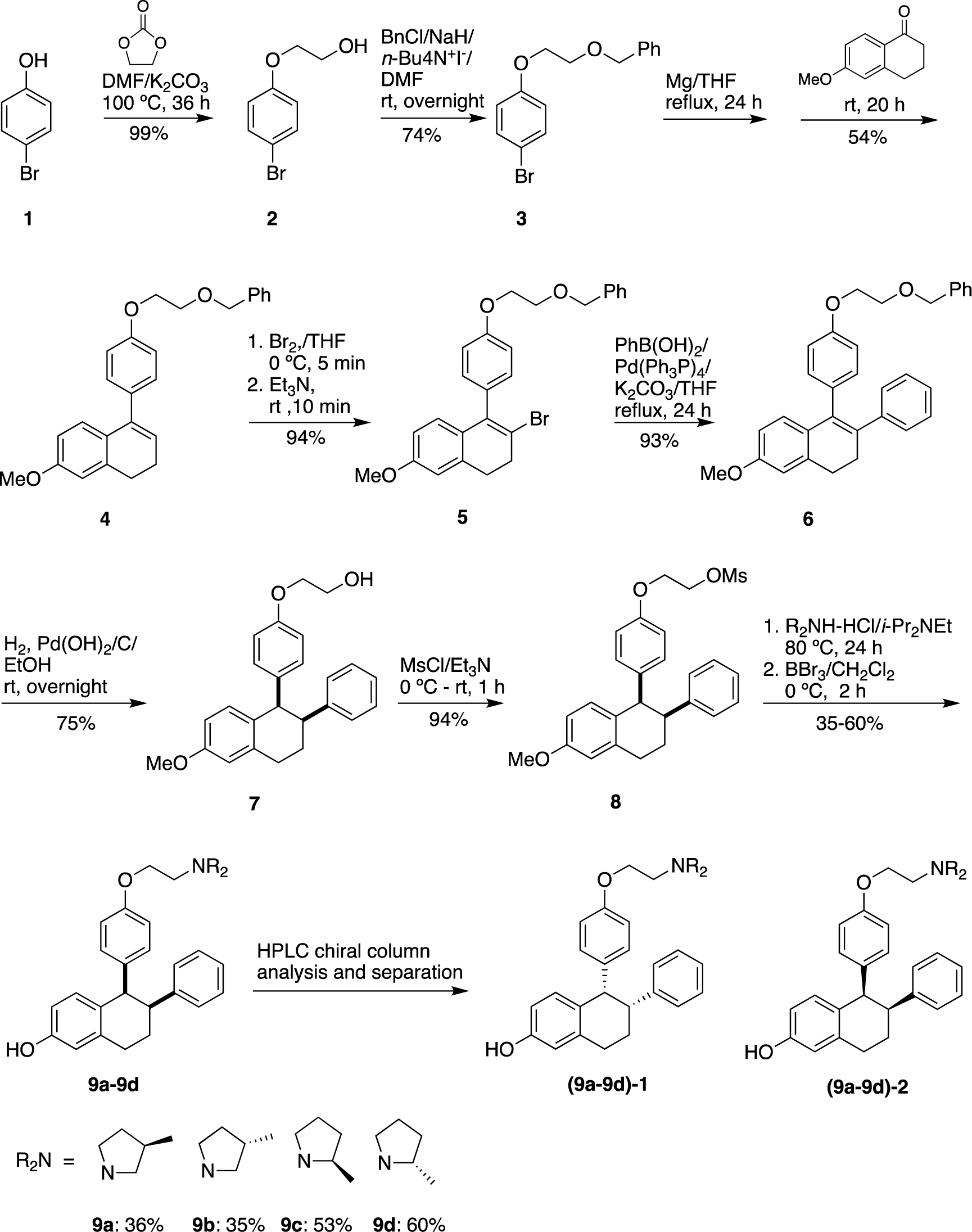
Experimental for the synthesis of Lasofoxifene analogues **(9a-9d)**.

**Figure 7-figure supplement 2:**
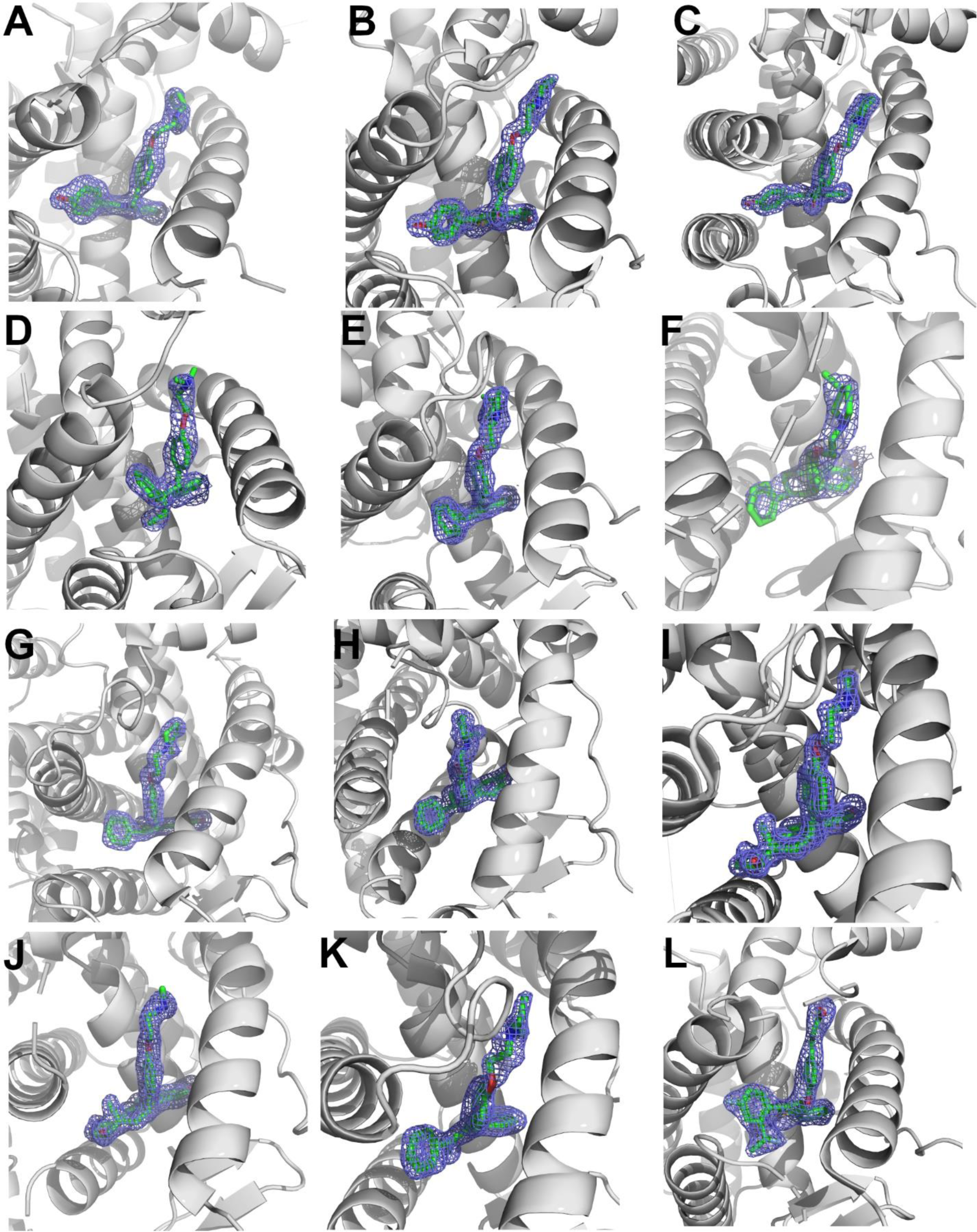
2mFo-DFc difference maps for antiestrogens in complex with WT and Y537S ERα LBD contoured to 1.5σ. A) Y537S-BZA. B) WT-RAL. C) Y537S-RAL. D) Y537S-4OHT. E) WT-LA-Deg. F) Y537S-LA-Deg. G) WT-LA-Stab. H) Y537S-LA-Stab. I) WT-RU39411. J) Y537S-RU39411. K) WT-Clomiphene. L) Y537S-LSZ102. All ligands are shown as sticks, all maps are shown as blue mesh, and all protein are shown as light grey ribbons.

**Figure 7-supplemental file 2:**
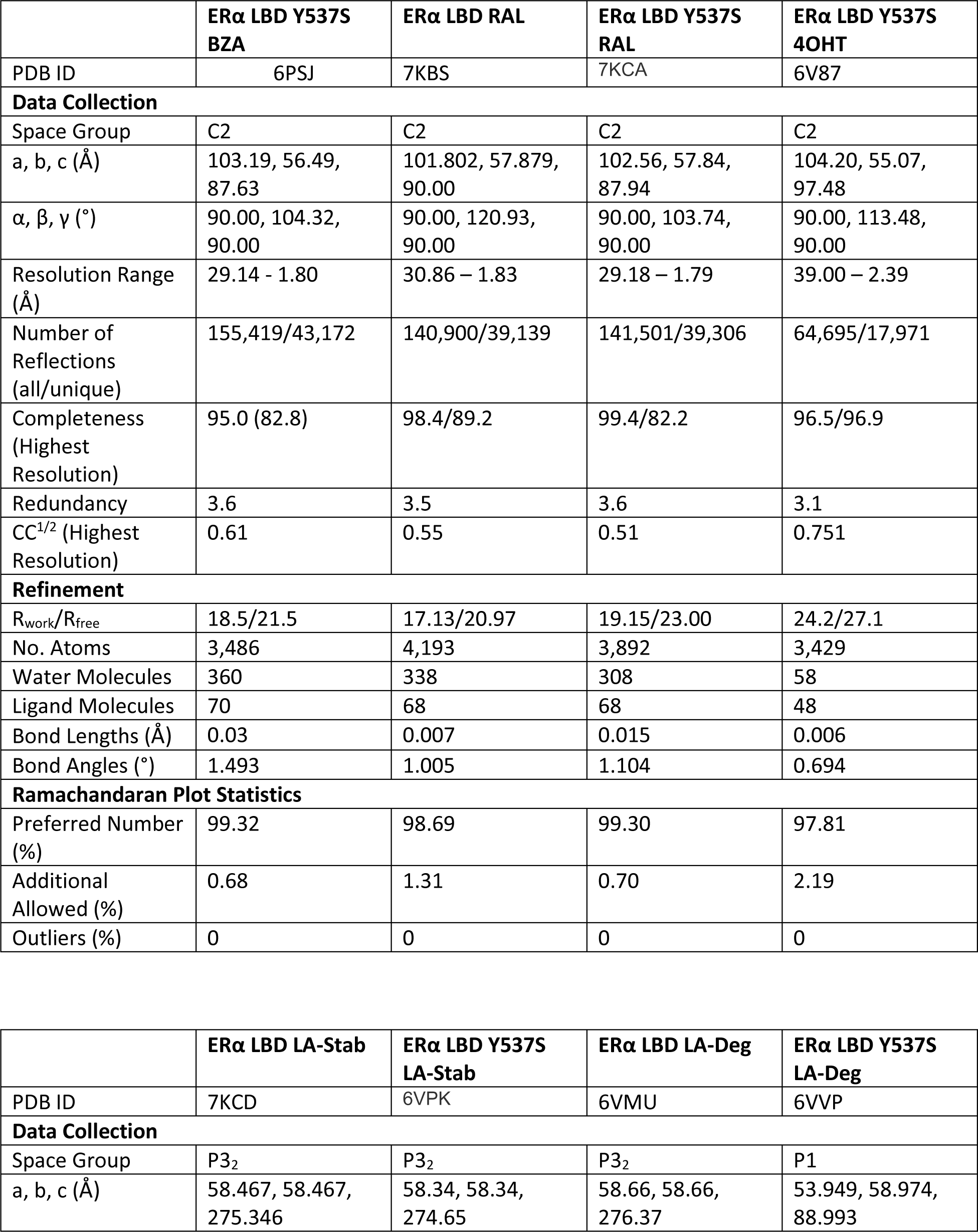
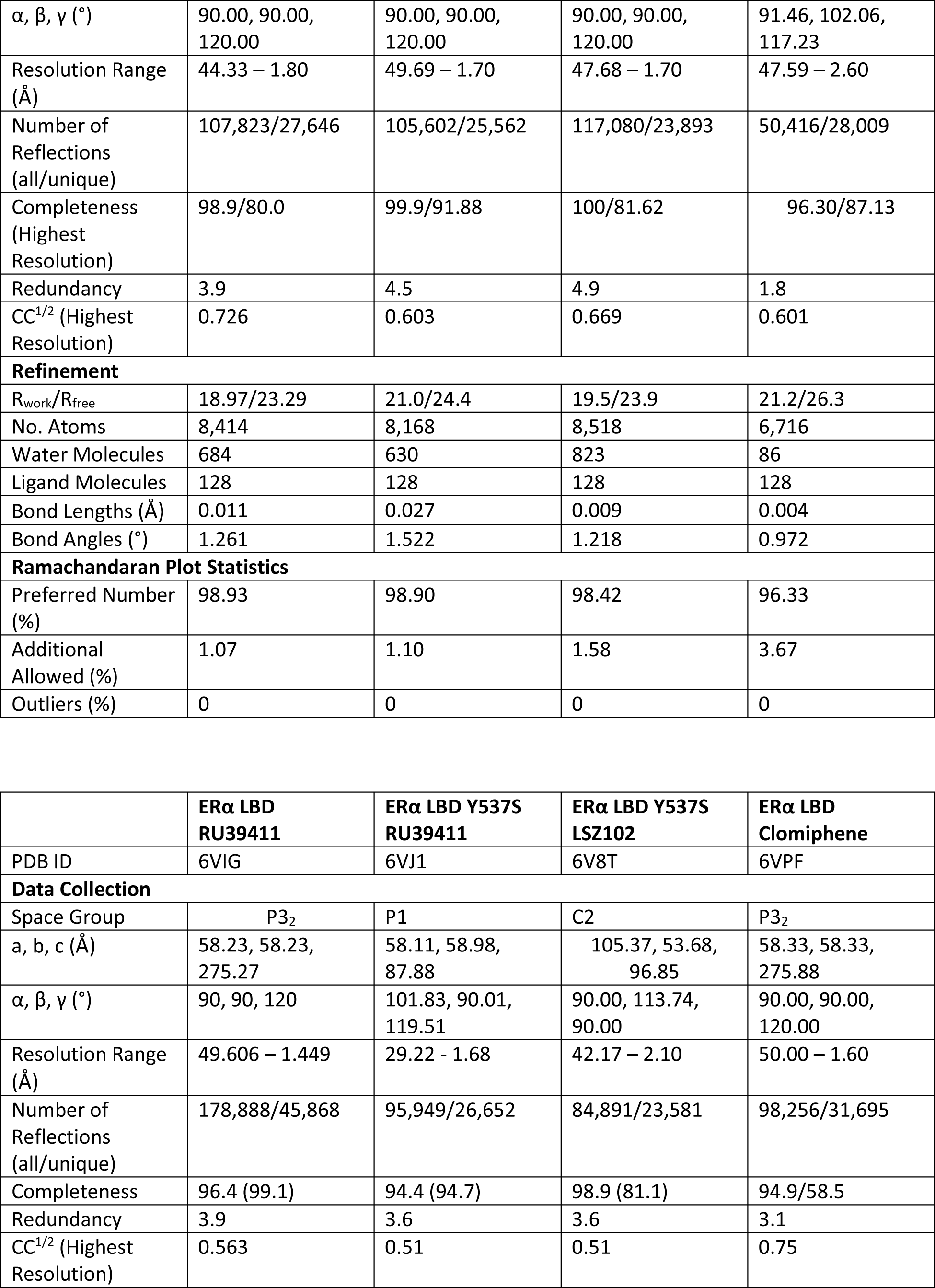
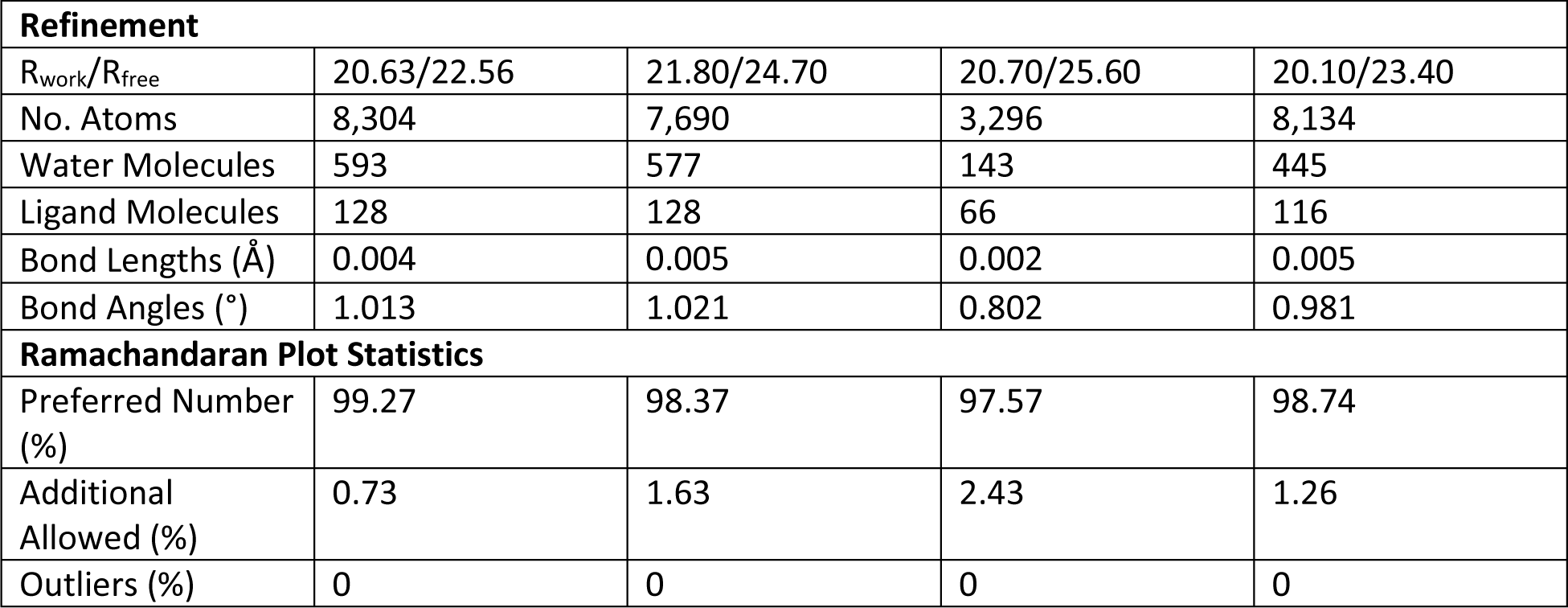
X-ray Crystal Structure Data Collection and Refinement

